# True-to-scale DNA-density maps correlate with major accessibility differences between active and inactive chromatin

**DOI:** 10.1101/2022.03.23.485308

**Authors:** Márton Gelléri, Shih-Ya Chen, Aleksander Szczurek, Barbara Hübner, Michael Sterr, Jan Neumann, Ole Kröger, Filip Sadlo, Jorg Imhoff, Yolanda Markaki, Michael J. Hendzel, Marion Cremer, Thomas Cremer, Hilmar Strickfaden, Christoph Cremer

**Affiliations:** Institute of Molecular Biology (IMB), Mainz, Germany; Biocenter, Dept Biology II, Ludwig Maximilian University (LMU), Munich, Germany); Max Planck Institute for Chemistry, Mainz, Germany; Interdisciplinary Center for Scientific Computing (IWR), University Heidelberg, Heidelberg, Germany; Neuroconsult GmbH, Heidelberg, Germany; Department of Oncology, Cross Cancer Institute, University of Alberta, Edmonton, Alberta, Canada; Kirchhoff Institute of Physics, University Heidelberg, Heidelberg, Germany

**Keywords:** active and inactive nuclear compartments (ANC and INC), true-to-scale DNA density maps (Mbp/µm^3^), microinjected nanobeads, replication domains (RDs), single molecule localization microscopy (SMLM), structured illumination microscopy (SIM), electron spectroscopic imaging (ESI), quantitative image analysis, Voronoi-tessellation

## Abstract

Chromatin compaction differences may have a strong impact on accessibility of individual macromolecules and macromolecular assemblies to their DNA target sites. Estimates based on fluorescence microscopy with conventional resolution, however, suggested only modest compaction differences (∼2-10x) between active and inactive nuclear compartments (ANC and INC). Here, we present maps of nuclear landscapes with true-to-scale DNA-densities, ranging from <5 Mbp/µm^3^ to >300 Mbp/µm^3^. Maps were generated from individual human and mouse cell nuclei with single-molecule localization microscopy at ∼20 nm lateral and ∼100 nm axial resolution and supplemented by electron spectroscopic imaging. Microinjection of fluorescent nanobeads with sizes corresponding to macromolecular assemblies for transcription and replication into nuclei of living cells, demonstrated their localization and movements within the ANC and exclusion from the INC.

## Introduction

Understanding the functional behavior of genomes requires knowledge of the dynamic biochemical and physical chromatin loop organization (Gasser and Laemmli, 1987; Neguembor et al., 2021) and its consequences for the accessibility of individual macromolecules, condensates with proteins and RNAs, as well as of structured macromolecular assemblies involved in nuclear functions. Microscopic studies of nuclear architecture have provided evidence for a multitude of chromatin ensembles with chromatin domains (CDs), chromosome territories (CTs), as well as active and inactive nuclear compartments (Cremer et al., 2020a; Cremer et al., 2015; Maslova and Krasikova, 2021; Nguyen et al., 2020; Nir et al., 2018; Strickfaden, 2021). Chromatin ensembles, such as topologically associating domains (TADs) and A- and B-compartments, were independently identified by genome wide measurements of 3D chromatin contacts (Dekker et al., 2017; Di Stefano et al., 2016; Kempfer and Pombo, 2020; Moretti et al., 2020) (for a glossary of currently used nomenclatures and conceptual differences, see supplemental information in (Cremer et al., 2020b). Physical maps of nuclear landscapes, supplemented with biochemical information about epigenetic markers and attachment sites of specific macromolecules and functional complexes, have revolutionized our views on nuclear organization. How nuclear compartments are established and maintained, and how they interact functionally, has remained a largely unsolved problem (Belmont, 2021; Misteli, 2020). Higher order chromatin arrangements differ substantially not only between different cell types and species, but also from cell to cell of a given cell population of the same cell type (Cardozo Gizzi, 2021; Strickfaden, 2021). This variability may reflect stochastic features of 4D spatial genomics (Finn and Misteli, 2019), while other features, including the space-time organization of nuclear landscapes at mesoscale, reflect a functionally important preciseness of genome organization (see Discussion).

In this study, we deal with the still unresolved problem of true-to-scale compaction levels (in terms of Mbp/µm^3^) of chromatin ensembles, constituting active and inactive nuclear compartments. These were previously identified by topological linking of relative chromatin density maps with functionally relevant nuclear hallmarks (Cremer et al., 2020a; Cremer et al., 2015) or by 3D-DNA contact frequency data (Lieberman-Aiden et al., 2009). At face value, a large body of evidence obtained in previous studies suggests only modest compaction and accessibility differences for functional macromolecules and macromolecular aggregates between active and inactive chromatin ensembles. Our present study challenges this concept.

Figure 1a-b shows views of higher order chromatin organization based on apparently ‘open’ chromatin loop configurations. Accordingly, an interchromatin space expanding both within the interior and outside of individual loops seems wide enough to allow penetration of individual macromolecules and complete macromolecular machineries (Figure 1c). In contrast, concepts of a much more ‘crumpled’ organization of chromatin (Figure 1d), and a ‘closed’ configuration of densely packaged nucleosomes (Figure 1e), may constrain the penetration of individual macromolecules into the interior of chromatin ensembles, and exclude the penetration of macromolecular machineries or phase-separated condensates (Maeshima et al., 2015). Our current version of the ANC-INC model (Figure 1f) supports this concept (Cremer et al., 2020a) (see below).

**Figure 1:**
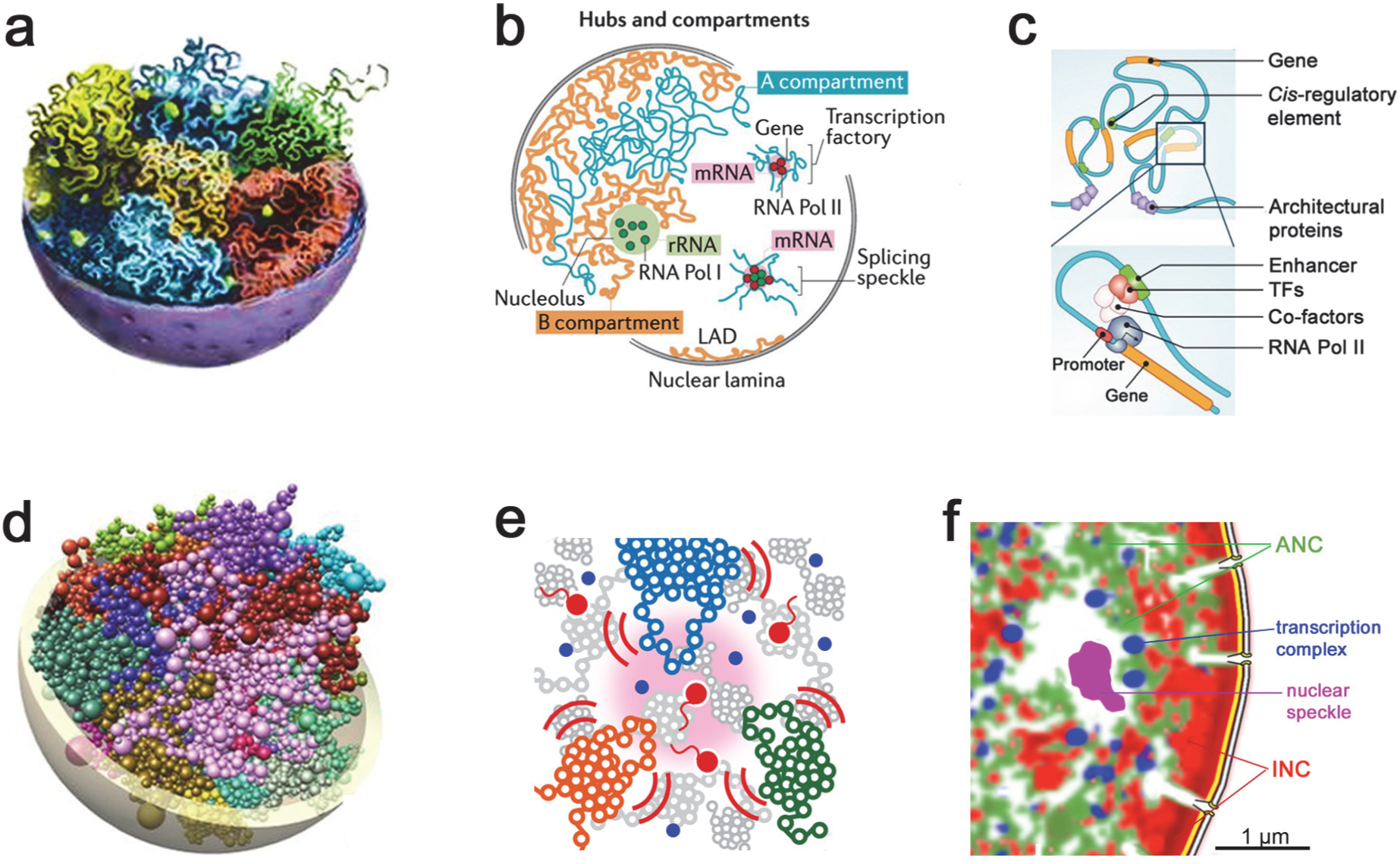
Views of 3D nuclear landscape. **a** Voronoi-tessellated SMLM-fBALM image of a mid-optical section (thickness ∼100 nm through a diploid human fibroblast cell nucleus (BJ1) stained with Sytox Orange (see Methods for further explanation how this image was generated). The color code indicates the estimated range of 3D DNA densities from <5 Mbp/µm^3^ to >40 Mbp/µm^3^, reflecting 2D SMP densities <2,500 SMPs/µm^2^ to >20,000 SMPs/µm^2^ (see Supplemental Figure S3, Supplemental Videos S1a,b) and Results for 3D DNA densities estimated from 2D SMP densities in 100 nm thick sections). **b, 1-3** Magnifications of the three boxed areas in (a) demonstrate sites located at the nuclear periphery (1) and in the nuclear interior (2, 3); edge length 3000 nm. Asterisks denote IC-lacunae. **b, 1’–3’** Further magnification of boxed areas in (1-3), edge lengths 1000 nm (1’), and 500 nm (2’,3’). **c, 1’’–3’’** upper row: Further magnification of boxed areas in (1’-3’), edge length 200 nm. lower row: Individual Voronoi centers (seeds) are shown for the same Voronoi areas as in 1’’ – 3’’. **d, 1’’’–3’’’** same areas as in c 1’’ depicted here with another color code (shown at bottom) for 3D DNA densities from 40 Mbp/µm^3^ to about 350 Mbp/µm^3^, reflecting a range of 2D SMP densities between ∼20,000 and ∼180,000 SMPs/µm^2^.

It is of note that many published cartoons show details of nuclear landscapes that are not drawn to scale. True-to-scale maps of spatial chromatin compaction, however, is a fundamental issue of chromatin accessibility and inaccessibility with major functional implications (see Discussion). For a comprehensive solution of this issue, super-resolved maps across entire nuclear landscapes are required, providing absolute local DNA densities (Mbp/µm^3^) instead of relative density differences (Schmid et al., 2017). Studies performed with the resolution limit of conventional fluorescence microscopy (∼200 nm in the object plane, and ∼600 nm along the optical axis), have suggested relatively small density differences between euchromatin and heterochromatin regions (Gorisch et al., 2003; Lenart et al., 2003; Verschure et al., 2003). Insufficient resolution, however, underestimates the true density related differences between euchromatic and heterochromatic regions, comparable to a landscape of a mountain region, recorded from an airplane with a poor resolution camera, that levels the differences between high mountains and deep valleys. Estimates of absolute nucleosome densities in live HeLa cell nuclei, obtained by a combination of Fluorescence Correlation Spectroscopy and Confocal Imaging (Weidemann et al., 2003), revealed about tenfold intranuclear differences in local nucleosome density, ranging from about 25 µM to 250 µM. Assuming 200 bp per nucleosome (including linker DNA), these values amount to about 3 Mbp/µm^3^ for low density regions and 30 Mbp/µm^3^ for high density regions. The authors mentioned that they could not exclude the possibility of the formation of local barriers only detectable with a much higher structural resolution. Studies of nuclei in live-murine cells performed with laser confocal microscopy (Imai et al., 2017) suggested a moderate accessibility barrier of non-nucleosomal materials (proteins/RNAs) that constrains the diffusion of proteins into heterochromatin.

Whereas studies with conventional optical resolution comprises an observation volume in the order of ∼10,000 nucleosomes (Weidemann et al., 2003), the improved structural resolution obtained with single molecule localization microscopy (SMLM) (for reviews see (Cremer and Masters, 2013; Gould and Bewersdorf, 2012; Khater et al., 2020; Lelek et al., 2021; Schermelleh et al., 2019) allows the visualization of individual clusters with less than 100 nucleosomes. Previous microscopic studies performed at this highly enhanced resolution level have indicated dense assemblies of nucleosomes in heterogeneous groups comprising a few kb of DNA, termed nucleosome clutches (NCs) (Ricci et al., 2015). NCs provide candidate structures that may represent building blocks for all higher-order nucleosome structures above the single nucleosome level, such as chromatin domains or TAD-like structures (Bintu et al., 2018; Ricci et al., 2015; Szabo et al., 2019; Szabo et al., 2018).

The present quantitative, high-resolution study presents the first true-to-scale 3D DNA density maps of the absolute heights of DNA density peaks and valleys across entire nuclear landscapes of individual cells in terms of Mbp/µm^3^ at nanometer-scale resolution. Differences between high density chromatin structures assigned to the INC and low DNA density regions assigned to the ANC (Babokhov et al., 2020; Cremer et al., 2020a) range from <5 Mbp/µm^3^ to >300 Mbp/µm^3^ and are much larger than DNA compaction differences claimed in previous studies with conventional microscopic resolution. In addition, we performed microinjection experiments with 20 and 40 nm sized fluorescent nanobeads, reflecting sizes of macromolecular assemblies for transcription and replication (Maeshima et al., 2015), and demonstrate their preferential enrichment in the active nuclear compartment (ANC) and exclusion from the inactive compartment (INC). These findings support a substantially lower compaction in the ANC associated with a higher accessibility of chromatin ensembles compared to the INC, with major implications for the functional organization of the cell nucleus.

## Results

### Determination of absolute DNA densities across nuclear landscapes

Single Molecule Localization Microscopy (SMLM) (Cremer et al., 2017) combined with fluctuation assisted Binding Activated Localization Microscopy (fBALM) (Szczurek et al., 2018; Szczurek et al., 2017) was used to record millions of blinking events of the fluorophore Sytox Orange, bound to double-stranded DNA in single light optical nuclear sections from morphologically intact human and mouse fibroblasts and from HeLa, as an example of a human cancer cell-line. Since Syto Nucleic Acid Stains (ThermoFisher Scientific) bind to both DNA and RNA, RNA was removed by treatment with an RNase cocktail before staining with Sytox Orange. In line with our major goal to determine absolute DNA densities of euchromatic and heterochromatic regions at high resolution, Sytox orange shows a more equal affinity to both GC and AT rich DNA, compared to DAPI and Hoechst 33342, which bind preferentially to AT rich DNA (Chazotte, 2011) (Supplemental Figure S1).

Figure 2 presents the color-coded, Voronoi-tessellated true-to-scale 3D DNA density SMLM-fBALM image obtained from a mid-plane optical section through a human fibroblast nucleus (BJ1) (Figure 2a) in G1 stage (Supplemental Figure S2). The image was generated from ∼1.5 million spatially mapped blinking events, registered in 50,000 consecutively recorded frames (Supplemental Figure S3). Single Molecule Positions (SMPs) of these events were located within an optical section of about 100 nm thickness with an average localization precision of ∼50 nm along the optical axis and an average lateral localization precision of ca. 8 nm (for details of this procedure see Methods). Figures 2b, c (upper panels) show selected regions with increasing magnifications. For panels a-c, a color-coded scale bar for 3D DNA densities with emphasis on the range >5 Mbp/µm^3^ and <40 Mbp/µm^3^ is presented at the top of Figure 2b. Figure 2c (lower panel) presents the Voronoi tessellations used for reconstructions of images 2c (upper panel). Figure 2d presents the same panel shown in Figure 2c with another color-bar code (below) that emphasizes the range of 3D DNA densities from >40 Mbp/µm^3^ up to ∼350 Mbp/µm^3^.

**Figure 2:**
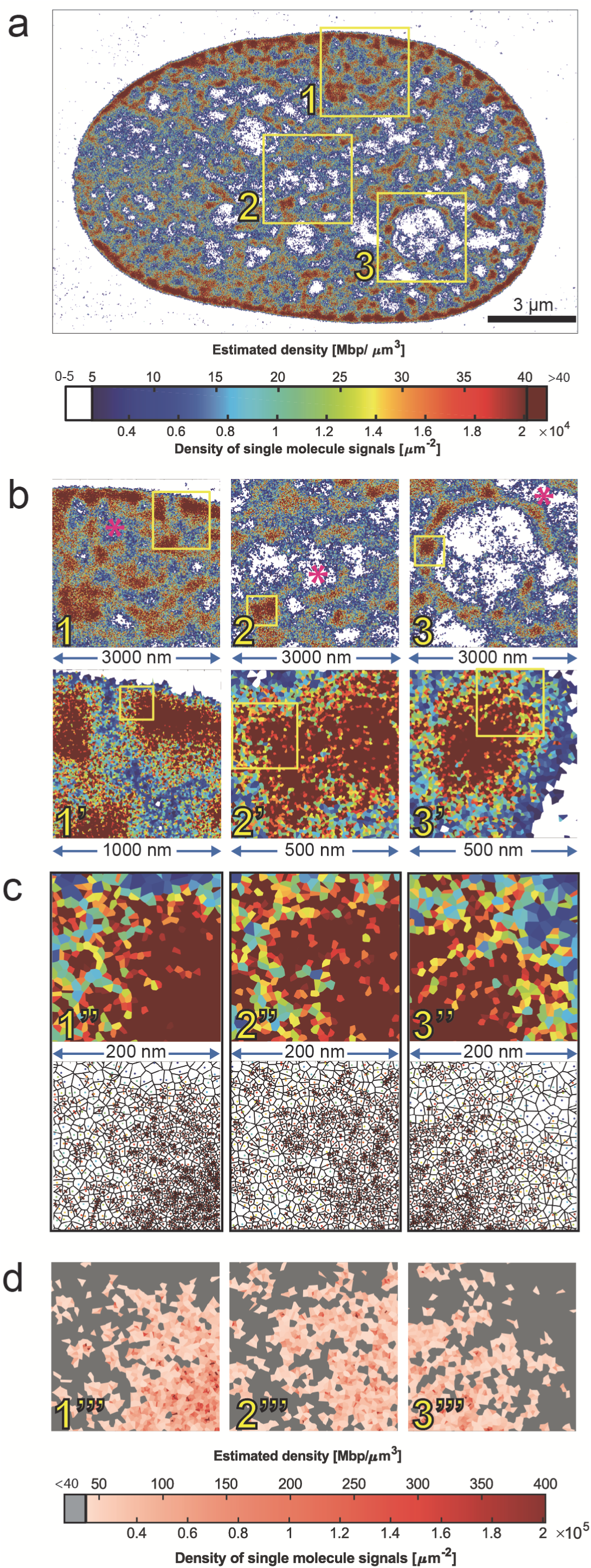
High-resolution, true-to-scale DNA density landscape of a human fibroblast nucleus. Current views of nuclear organization at mesoscale agree that chromosomes occupy distinct territories (CTs)/domains (Cremer and Cremer, 2010; Cremer et al., 2015; Dekker et al., 2017; Di Stefano et al., 2016; Kempfer and Pombo, 2020; Moretti et al., 2020; Paulsen et al., 2018). Current concepts of chromatin compaction, however, range from relatively open chromatin loop configurations **a** (Dekker et al., 2017); **b** (Kempfer and Pombo, 2020); **c** (Moretti et al., 2020) to proposals as in **d** describing the crumpled organization of chromatin as little balls without giving an opinion on the density of chromatin inside (Paulsen et al., 2018). **e** Higher order chromatin organization of chromatin based on nucleosome clusters (NCs)(Babokhov et al., 2020). Red dots: RNAPII; blue dots: transcription factors. NCs may form the first level of higher order chromatin structures beyond individual nucleosomes (Ricci et al., 2015). **f** ANC-INC model of nuclear architecture (Cremer et al., 2020a; Cremer et al., 2015). The active nuclear compartment (ANC) comprises the Interchromatin compartment (IC). IC-channels start at nuclear pores. The IC forms a network throughout the nuclear interior and expands at certain sites into wider IC-lacunae, which carry splicing speckles and other nuclear bodies. It is lined by the perichromatin region (PR) with less dense and more accessible chromatin (green). The inactive nuclear compartment (INC) carries more compact and poorly accessible chromatin (red); for further details see Figure 6 and text.

Absolute DNA densities (Mbp/µm^3^) were calculated as follows: In addition to the total number of SMPs, we estimated both the volume, comprising about 1/20th of the nuclear volume, and the total DNA content of the 100 nm thick optical section. From these numbers, we calculated that a single SMP in this BJ1 nucleus was representative on average for about 200 bp, that is roughly the DNA surrounding a single nucleosome (see Methods for further details). Next, we used 2D Voronoi-tessellation (Andronov et al., 2016; Levet et al., 2015) for partitioning 2D SMP density images into 2D polygonal images. Assuming that each 2D polygon represents in fact a 3D polygonal prism with a depth of 100 nm, we calculated the average DNA density for each prism by dividing its DNA content (based on the number of SMPs) by its volume. Polygonal Voronoi prisms were color-coded corresponding to 3D DNA density estimates, ranging from <5 Mbp/µm^3^ (white) to >40 Mbp/µm^3^ (deep red), respectively, as indicated in the color-coded scale for 2a-c. A further unfolding of color codes for densities >40 Mbp/µm^3^ up to ∼350 Mbp/µm^3^ and corresponding SMP densities is provided in the color scale column in Figure 2d.

Figure 3 exemplifies a mid-optical section of a nucleus from the near triploid human cancer cell line HeLa. Based on its small size, we assume that this nucleus also represents a G1 stage (Cremer et al., 2003). This color-coded, Voronoi-tessellated SMLM-fBALM image was constructed from about 1.2 million SMPs of DNA-bound, blinking Sytox Orange molecules registered in 50,000 consecutive frames within a 100 nm thick optical section. True-to-scale DNA densities were estimated as described above. Note that estimates of absolute DNA densities estimates are conservative and would increase by a factor 2 in case of a G2 DNA content. Towards the nuclear periphery, we note high-density chromatin ensembles with diameters up to several hundred nm, surrounded by chromatin with low DNA densities. Supplemental Figure S4 presents the SMLM-fBALM image of the mid-optical section from another HeLa cell nucleus, which shows the typical structural features of the ANC-INC model, but differs substantially from the example shown in Figure 3 with regard to a more expanded ANC at the cost of a reduced INC. Arguably, this nuclear landscape may reflect a more advanced interphase stage (Nagano et al., 2017). A detailed study of different interphase stages, however, was beyond the scope of our present study.

**Figure 3:**
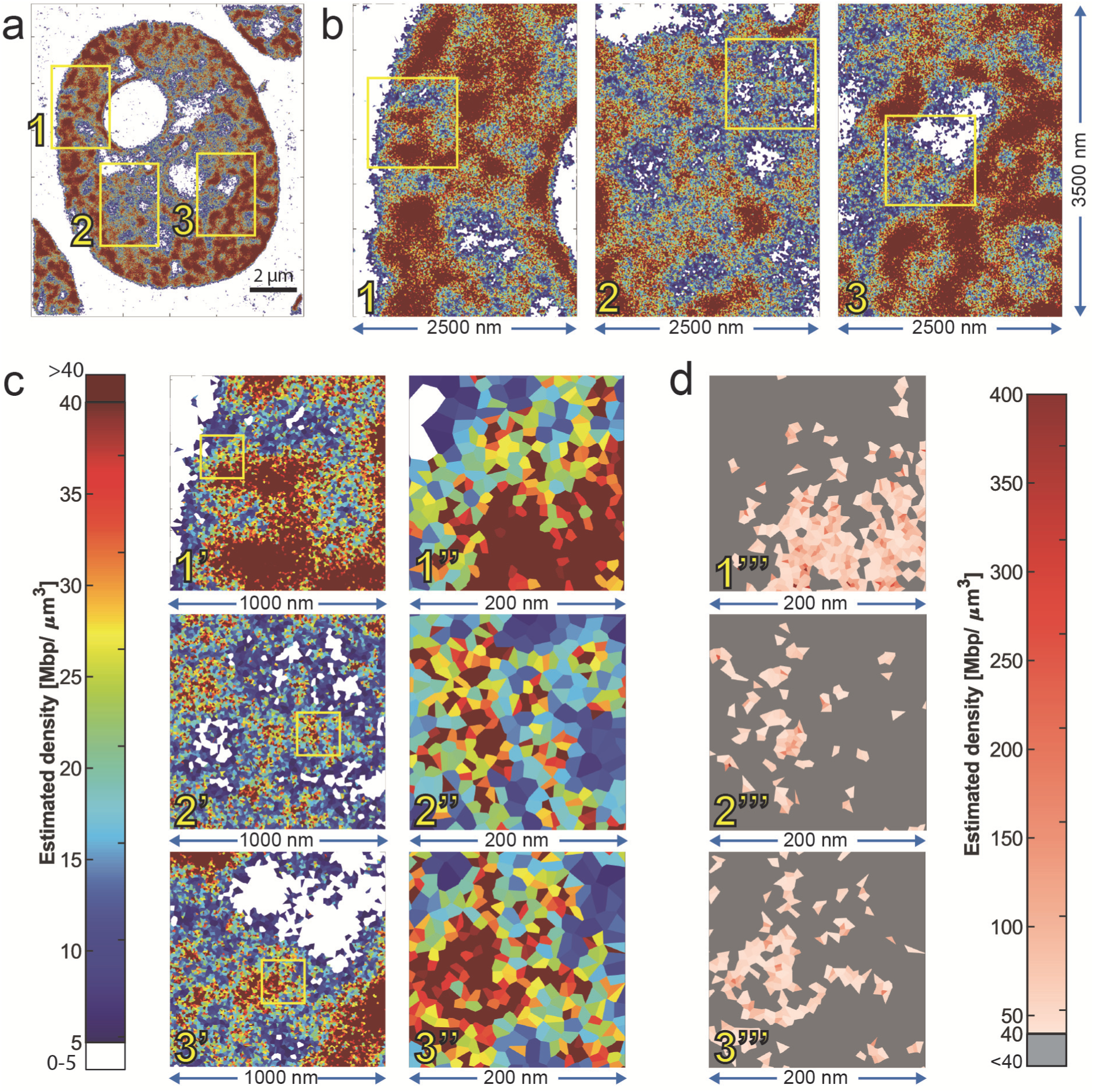
High-resolution, true-to-scale DNA density landscape of a HeLa cell nucleus. **a** SMLM-fBALM image (Voronoi-tessellation) of a mid-optical section (thickness ∼100 nm) through a HeLa cell nucleus. **b, 1-3** Magnifications of the three boxed areas in (a); edge lengths 2.5 µm x 3.5 µm. **c** color code shown on the left presents estimates of absolute densities for 0-5 Mbp/µm^3^ (white), and the range from 5 to (dark blue) to >40 Mbp/µm^3^ (dark red) (as described in Figure 2). **c, 1’-3’** Further magnification of boxed areas in (b,1-3); edge length 1000 nm; **c, 1’’-3’’** Further magnification of boxed areas in c,1’- 3’; edge length 200 nm, depicts individual color-coded Voronoi cells and clusters with the same color. **d, 1’’’- 3’’’** presents the same regions as in c 1’’- 3’’ shown in another color code (also described in Figure 2) to demonstrate the range of apparent DNA densities from 40 Mbp/µm^3^ (light red) up to maximum densities of 400 Mbp/µm^3^ (dark red).

In summary, Figures 2 and 3 exemplify the potential of our approach to identify the widely different, true-to-scale 3D DNA densities of chromatin ensembles and their nuclear arrangements across entire nuclear landscapes at mesoscale and nanoscale structural resolution.

Figure 4 shows a comparison of the DNA landscapes of a mouse C3H 10T1/2 fibroblast nucleus obtained from a Voronoi-tessellated SMLM-fBALM image (Figure 4a) and from electron spectroscopic imaging (ESI) (Hendzel and Bazett-Jones, 1996) (Figure 4b-e) with the same thickness (100 nm) of light optical and physical ultra-thin midsections, respectively. The two mouse fibroblast nuclei vary with regard to the more expanded IC (blue) in the nucleus studied with ESI; apart from this their DNA-density landscapes appear remarkably similar with less dense chromatin in the PR. In both human and mouse nuclei we note an enrichment of highly compact DNA at the nuclear periphery. In contrast to the human fibroblast nucleus, the mouse fibroblast nucleus carries prominent clusters of constitutive heterochromatin, called chromocenters (depicted by arrows).

**Figure 4:**
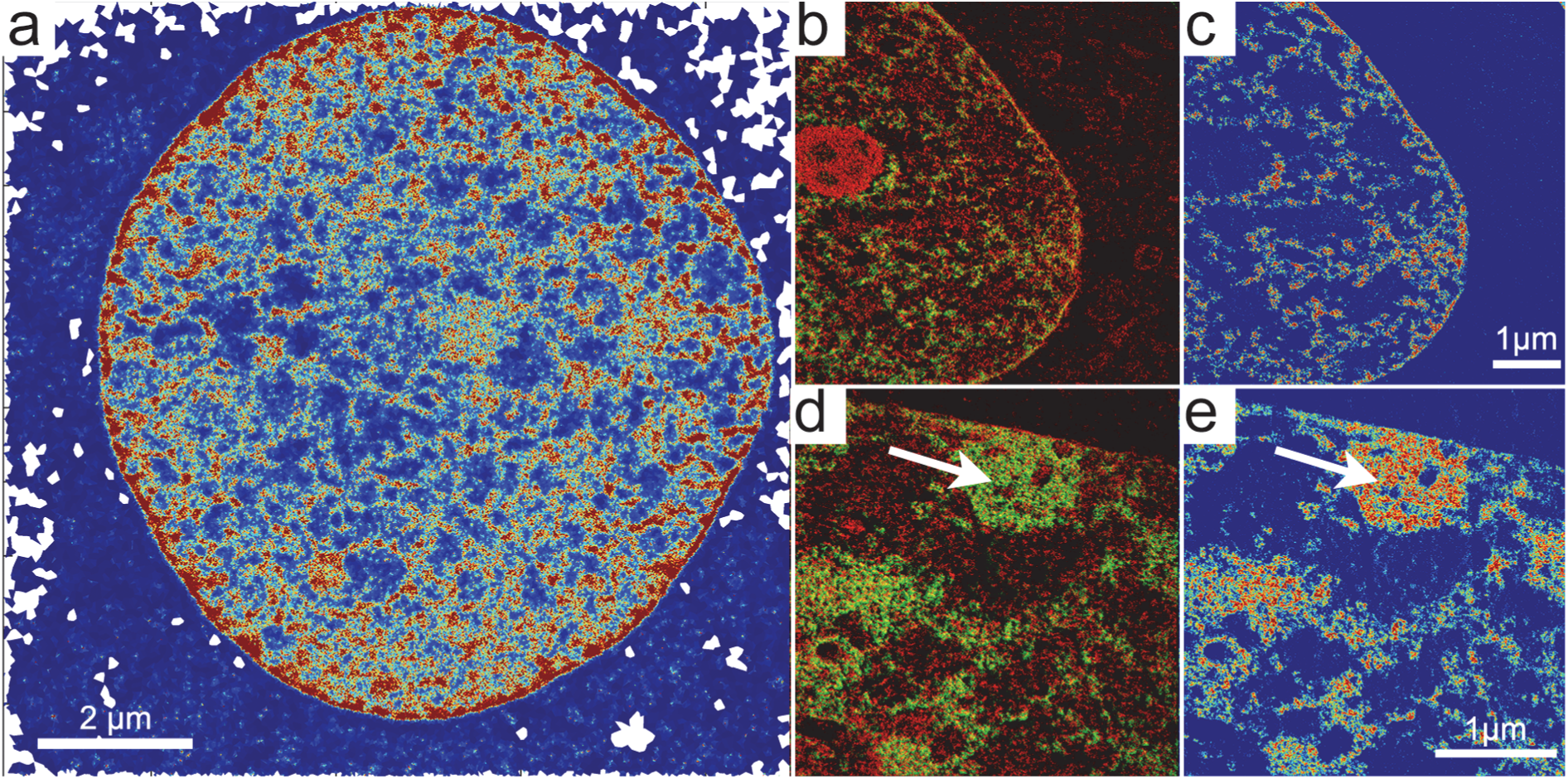
Comparison of high-resolution DNA landscapes of mouse C3H T1/2 fibroblast nuclei recorded with SMLM and ESI. **a** 100 nm optical mid-section through a mouse C3H T1/2 fibroblast nucleus recorded with SMLM-fBALM. **b-e** ESI micrographs taken from acrylic 100 nm thin mid-sections through another C3H T1/2 fibroblast nucleus. **b** and **d** show superimposed elemental maps of phosphorus (green) and nitrogen (red). **c** and **e** present the phosphorus map only, color-coded for different phosphorus densities as a reflection of DNA densities. For better comparison, the same color-coded look-up table (jet) was used for SMLM (a) and ESI images (c,e; blue indicates lowest DNA densities that belong to the IC, lined by light blue and green colored intermediate densities (PR); red marks high DNA compaction attributed to the INC (compare Figures 2 and 3). Arrows in (d) and (e) indicate examples of chromocenters. In order to secure that only the DNA landscape was recorded, RNA was digested in both nuclei prior to imaging.

### Localization and constrained movements of microinjected nanobeads within the ANC

Evidence presented above supports DNA density differences, ranging from <5 Mbp/µm^3^ in the ANC to >300 Mbp/µm^3^ in the INC. For a rough idea of a sterical accessibility barriers imposed by chromatin, we estimated 3D distances between the surfaces of nucleosomes, represented by 10 nm sized spheres uniformly arranged in 3D space in comparison to a spherical protein with 5 nm diameter, and a 20 nm sized condensate or macromolecular machinery. We assumed DNA fully packaged with nucleosomes at different DNA densities of 5, 30, 60 and 150 Mbp/µm^3^ (Figure 5a). At face value, this scheme favors the hypothesis of an accessibility barrier of dense nucleosome aggregates attributed to the INC. Apparently, there is little room left for different geometrical arrangements of nucleosome spheres at very high DNA densities. On the contrary, geometrical nucleosome arrangements at low densities (<5 Mbp/µm^3^) provide sufficient nucleosome free space for a variety of individual loop structures, which allow a high degree of accessibility for macromolecules and macromolecular complexes/condensates. Since the space-time dynamics of chromatin domains in nuclei of living cells is currently not well understood (see Discussion), it is not possible to conclude from the observed DNA density differences in fixed cell nuclei, to which extent chromatin in the INC with high compaction levels may constrain the accessibility of individual proteins and RNAs or fully exclude macromolecular machineries and condensates, assembled within the ANC. For an experimental test, we microinjected fluorescent beads with 20 nm and 40 nm diameter, respectively, into nuclei of living HeLa cells (Figure 5b-e) and of C3H 10T1/2 cells (Figure 5f-i). The sizes of these nanobeads were chosen to approximate sizes of transcription- and replication-machineries or phase-separated droplets. HeLa cells used for microinjection experiments were either kept in isotonic (290 mOsm) medium (Figure 5b), or in hypertonic (570 mOsm) medium (Figure 5c), resulting in the rapid induction (∼1 min) of hyper-condensed chromatin (HCC). When cells with HCC were re-incubated in medium with physiological osmolarity (290 mOsm), nuclei with normally condensed chromatin (NNC) were quickly restored, even after several NNC-HCC-NCC cycles (Albiez et al., 2006). For both states of chromatin compaction, analysis of nanobead positions in fixed cell nuclei demonstrated their presence in less condensed chromatin attributed to the ANC (Figure 5d, e). Corresponding DNA (DAPI) intensity profiles are shown by line scans through nuclei encompassing the site of a nanobead (Supplemental Figures S5 and S6). Most HeLa nuclei with nanobeads, including cells after a NCC-HCC-NCC cycle, were able to pass through mitosis and formed two daughter cells with apparently normal nuclei. Notably, live cell imaging demonstrated mitotic cells with nanobeads excluded from condensed mitotic chromosomes (data not shown, see (Hübner, 2020).

**Figure 5:**
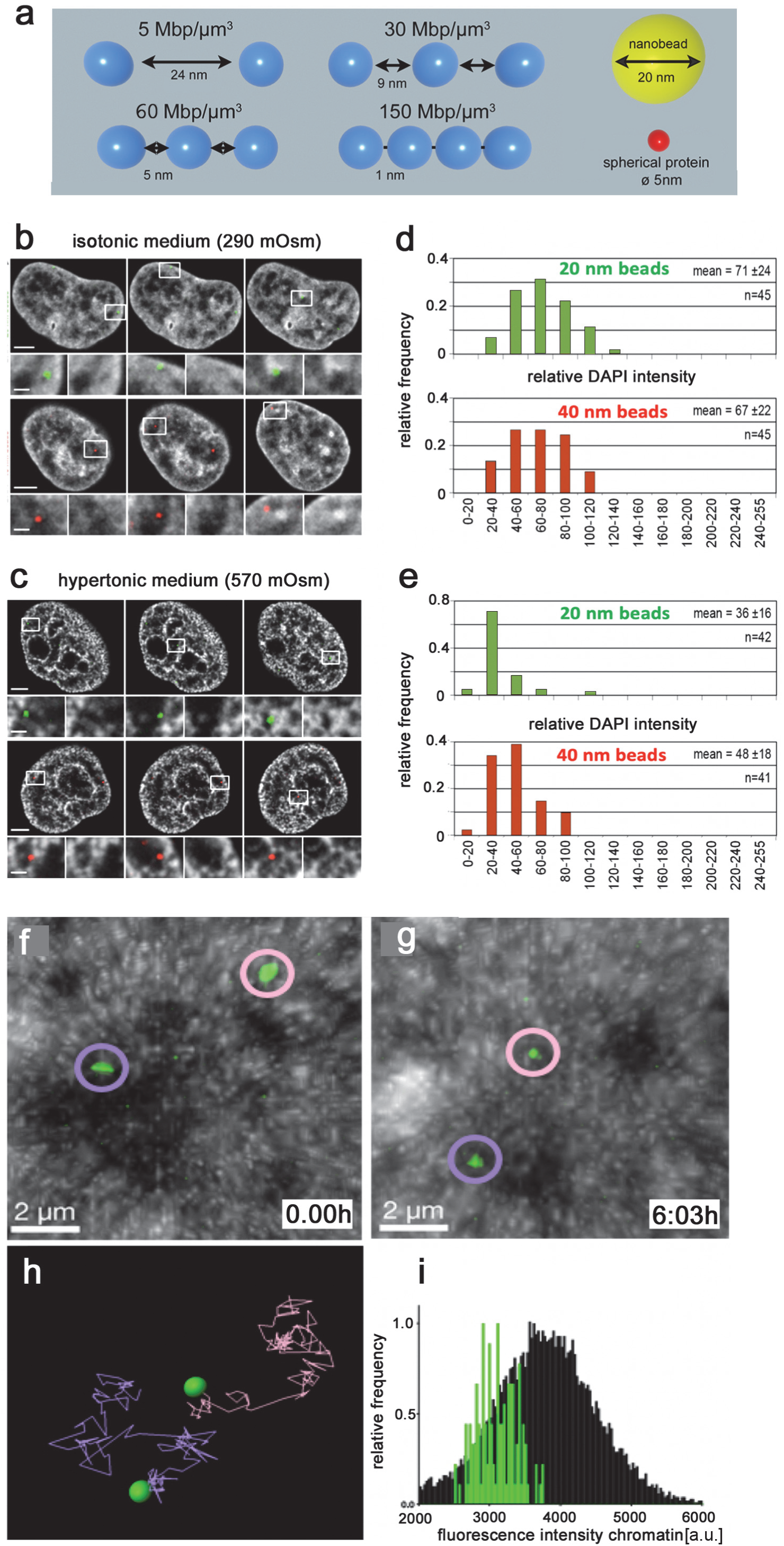
Localization and mobility of microinjected nanobeads within the ANC. **a** Distances between surfaces of uniformly arranged nucleosomes (spheres of 10 nm Ø) in 3D space estimated for different DNA densities (5, 30, 60, and 150 Mbp/µm^3^), compared to the size of individual macromolecules (Ø 5 nm) and of condensates or macromolecular complexes (Ø 20 nm) (for details see Methods). **b-c** Live HeLa cells, kept either in isotonic medium (**b**) or hypertonic medium (**c**), were microinjected with 20 nm (green, upper panel) and 40 nm (red, lower panel) fluorescent nanobeads and fixed thereafter with PFA. Nuclear DNA was counterstained with DAPI (gray). 3D confocal serial sections were recorded from microinjected cells, and sections with bright nanobead signals (boxed areas) were selected for further evaluation. Magnifications of the boxed areas (lower rows) show nanobeads located within the IC-lacunae (left) together with the DAPI image alone (right) for comparison. Intensity profiles of line scans through individual nanobeads and across the DAPI stained nuclear landscape were analyzed in 40 nuclei (Supplemental Figures S5 and S6). Scale bars: 4 μm in images of whole nuclei, 1 μm in magnifications. **d-e** Localization of 20 and 40 nm sized nanobeads as a function of relative DAPI intensities in nuclei of cells kept in isotonic (d) or hypertonic medium (e). In both isotonic and hypertonic nuclei, ≥90% of nanobeads are located in regions with DNA densities ≤100 (gray value range 0-256). Mean values are presented with standard deviation. These data demonstrate the location of nanobeads within less dense DNA attributed to the ANC. For further details see (Hübner, 2020). **f-g** In vivo labeling of the chromatin of C3H 10T1/2 cells was performed with a fluorescent protein (mRFP) associated with H2B. Dark and light regions show chromatin of low and high densities, attributed to the ANC and INC, respectively. The two green encircled dots indicate the position of two microinjected nanoparticles at the beginning and the end of the observation period. **h** shows the whole paths travelled by the two nanobeads. **i** A comparison of relative mRFP intensities (arbitrary units) across the entire nuclear section (black) with intensities at sites of nanobeads (green) supports the hypothesis that nanobeads move within less dense chromatin, attributed to the ANC (compare Supplemental Video S2).

In a second set of experiments, we studied movements of microinjected 20 nm sized fluorescent nanobeads in nuclei of live C3H 10T1/2 nuclei, where transient transfection of C3H 10T1/2 cells with a construct for the expression of histone H2B tagged with mRFP (Wolff et al., 2006) allowed the visualization of chromatin (Strickfaden et al., 2010). Figure 5f-g shows two nanobeads (green) within such a nucleus at the beginning and the end of a 6h observation period. Figure 5h presents the entire paths travelled by the two nanobeads; Figure 5i indicates that movements occurred most of the time within less dense chromatin (attributed to the ANC), possibly in a random-walk like fashion.

For a glimpse on dynamic topographical changes between co-aligned ANC and INC ensembles, we conducted DNA pulse labeling experiments during S-phase. These experiments indicate the formation of replication domains within the ANC. Pulse-chase labeling experiments demonstrate replication within wide IC-channels pervading constitutive heterochromatin blocks, followed by the re-location of replicated, inactive chromatin into dense chromatin attributed to the INC (Supplemental Figure S7).

## Discussion

In this study, we present examples of nuclear landscapes from a variety of human and mouse cell types imaged with different optical principles (SMLM, SIM and ESI). With SMLM-fBALM, the positions of millions of ‘blinking’ DNA-bound reporter fluorophore positions (SMPs) in optical nuclear sections were obtained with an average lateral resolution of ∼20 nm within the object plane and an axial resolution of ∼100 nm. Accordingly, the estimated observation volume (Hell and Stelzer, 1992) was about 500x smaller compared to conventional wide-field fluorescence microscopy. High-resolution DNA density maps generated from SMLM-fBALM images revealed absolute DNA compaction differences in nuclei of fixed cell samples from <5 Mbp/µm^3^ to >300 Mbp/µm^3^, a range much higher than previously published (see Introduction).

Monte-Carlo simulations (Maeshima et al., 2015) have predicted accessibility constraints for nucleosomal concentrations in the order of 0.3 mM to 0.5 mM, reflecting 40 – 60 Mbp/μm^3^ in case of DNA fully compacted with nucleosomes. Based on these simulations and our new results, we provisionally attribute chromatin with less dense DNA (<40 Mbp/µm^3^) to the active nuclear compartment (ANC), and chromatin with denser DNA (>40 Mbp/µm^3^) to the inactive nuclear compartment (INC). Constrained movements of individual macromolecules into the INC may help to increase the concentration of such molecules in the ANC and as a consequence lead to a preferential formation of functionally important macromolecular complexes in the ANC, composed of the interchromatin compartment (IC, and the perichromatin region (PR) (Cremer et al., 2020a). To explore sterical accessibility barriers imposed by densely packaged chromatin attributed to the inactive chromatin compartment (INC), we microinjected fluorescent beads with diameters of 20 nm and 40 nm, reflecting sizes typical for transcription- and replication-complexes, into nuclei of living cells. We observed that movements of these nanobeads were typically restricted to the less dense chromatin of the ANC. These results are consistent with previous live cell fluorescence microscopy of artificial nuclear bodies built up from clustered plasmids with inducible transgenes (Meggendorfer et al., 2010). Artificial bodies were formed within the IC and stayed there during interphase independently of their silent or induced active state, suggesting that their size prevented diffusion into the lining chromatin. Activation of transgene transcription resulted in the recruitment of RNA polymerase II and NFκB and a closer positioning to splicing speckles. Upon entry of cells into mitosis, the bodies were never trapped within, but always separated from mitotic chromosomes.

Our new observations together with published results support the hypothesis that functional machineries are size-excluded from the compact interior of chromatin ensembles with DNA densities allocated to the INC (Cremer et al., 2020a; Maeshima et al., 2015). The examples of nuclei shown in this study for a human fibroblast (BJ1), a human cancer cell (HeLa), and a mouse fibroblast (C3H T1/2), support typical architectural features of the ANC-INC model, arguably because these features are indispensable for functional reasons in cell nuclei. Many nucleosome clusters have sizes in the order of ∼100 nm or even less and therefore their spatial conformation cannot be further analyzed by microscopy at conventional resolution. In line with our present study, a series of previous microscopic studies with SMLM (Kirmes et al., 2015), and structured illumination microscopy (SIM) (Cremer et al., 2020b; Cremer et al., 2015; Miron et al., 2020) has provided evidence for major functional roles of chromatin ensembles located in the ANC, including transcription and DNA replication (Smeets et al., 2014). In addition to functionally active chromatin ensembles located in the perichromatin region (PR) (Rouquette et al., 2010), chromatin/DNA loops apparently extrude from the PR into the interchromatin compartment (IC). Arguably, large and highly expressed genes in normal cell types (Leidescher et al., 2022), as well as highly expressed amplicons of genes in cancer cells (Solovei et al., 2000), expand along IC-channels through the nuclear space into and even through neighboring CTs. These observations, however, do not contradict the hypothesis that loops formed by most active genes are small-sized. A schematic model of a chromatin domain (Figure 6a adapted from (Miron et al., 2020) argues for small loops with active histone marks (green dots), associated transcription factors and RNA polymerase complexes located at the CD periphery (ANC). In contrast, nucleosomes with repressive histone marks locate within the repressed CD core (INC), where they are inaccessible for the transcriptional machinery. This model implies that transcriptional repression or activation may depend on mechanisms for the 3D re-location and epigenetic remodeling of nucleosomes with important target sequences, such as from the transcriptionally competent CD periphery into the silent interior and vice versa from the interior to the periphery for activation. Notably, such a re-location mechanism allows stochastic heterogeneity of the positions of nucleosomes whose exact positions may be functionally irrelevant, but requires a functionally indispensable preciseness of the space-time organization of CDs only for those nucleosomes which carry important DNA target sequences, such as transcription regulatory elements (Figure 6b) (Cremer and Cremer, 2019). For example, we consider regulatory and coding sequences of active genes located within the ANC in cycling cells. Their relocation into the compact chromatin of the INC in postmitotic, terminally differentiated cells would provide a structural mechanism to safeguard long-term silencing. In case that regulatory sequences of such genes become inaccessible in the interior of highly compact domains, their accessibility would require chromatin decondensation to an extent that allows constrained movements of required transcription factors into the domain interior. Similar structural mechanisms may be envisaged for movements of DNA between neighboring parts of the ANC and INC required for DNA replication.

**Figure 6:**
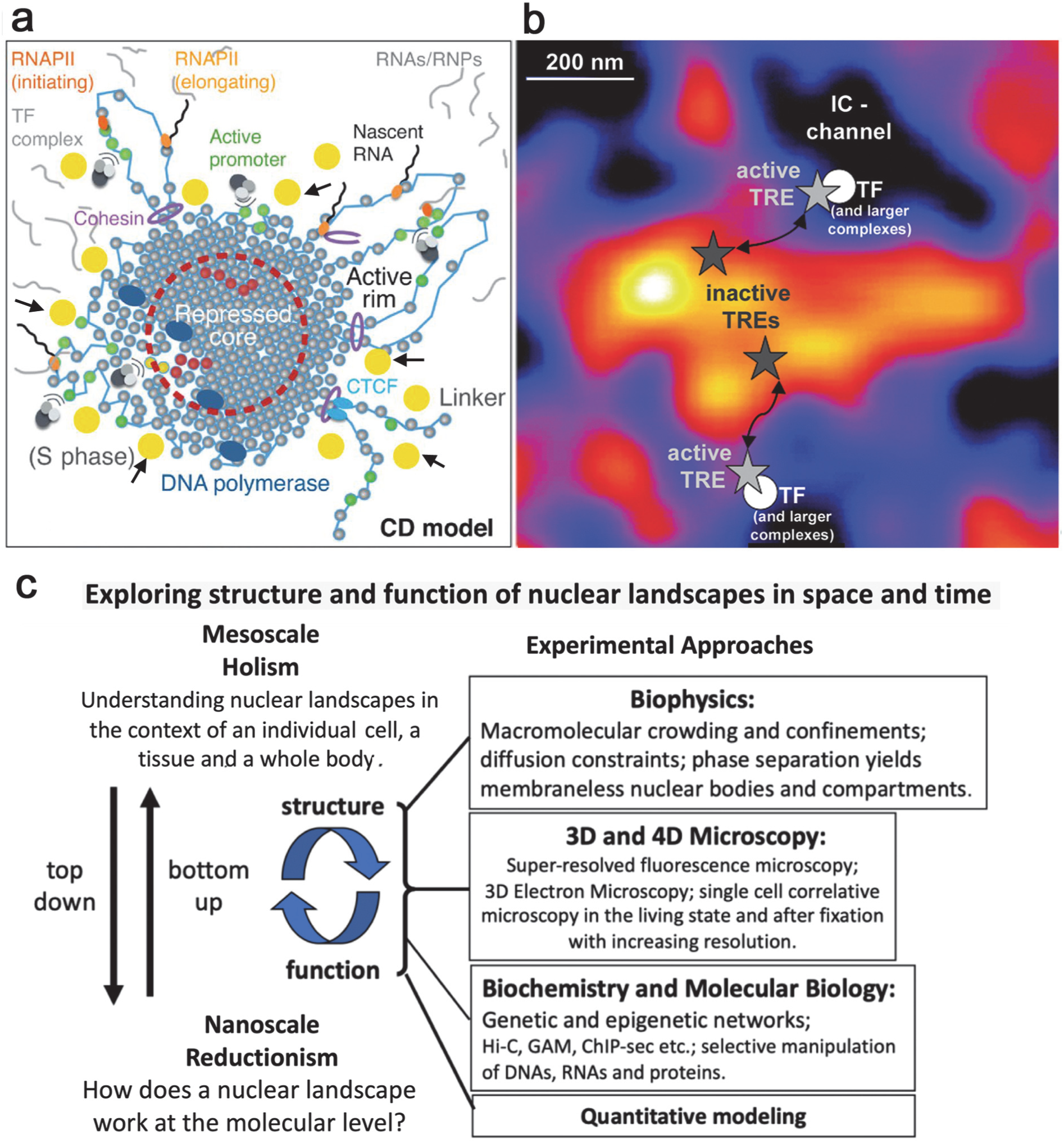
Nanoscale and mesoscale topography of the nuclear landscape. **a** Nanoscale model of a chromatin domain adapted from (Miron et al., 2020). It presents a chromatin domain with a repressed core of densely compacted nucleosomes and an active rim with expanding, transcribed chromatin loops. Yellow dots (some of them marked by small arrows) are added to indicate phase-separated, 20-nm sized droplets. **b** Mesoscale model of functionally interacting active and inactive nuclear compartments (Cremer and Cremer, 2019). The ANC-INC model proposes functional interactions between the ANC and INC based on hypothetical bidirectional chromatin movements, such as genes and corresponding transcription regulatory elements (TREs), from the ANC into the INC for long-term silencing, and from the INC to the ANC in case of renewed transcriptional activation. In this hypothetical nuclear section, the INC is represented by a chromatin domain cluster with several high-density peaks (white-yellow) and surrounding slopes (red), embedded within the ANC with its much less compacted chromatin. The ANC is composed of transversely cut IC-channels (black) and the lining perichromatin region (PR) (blue). Dark stars indicate the location of inactive TREs within the INC, light grey stars the position of active TREs in the ANC. **c** An integrative experimental approach to explore the intricate space-time, structure-function-conundrums of nuclear landscapes at nanoscale and mesoscale.

Still, it is not clear, how chromatin nanostructures form higher levels of chromatin organization (Beagrie et al., 2021; Eltsov et al., 2008; Jerkovic and Cavalli, 2021; Minnoye et al., 2021; Zakirov et al., 2022). For an in-depth understanding of these relationships, it is mandatory to produce fully integrated, high-resolution maps, based on the full range of microscopic/biophysical and biochemical approaches (Figure 6c). Recent evidence has suggested a dual nature of chromatin (Hansen et al., 2021; Strickfaden et al., 2020). The nuclear matrix proteome (Sureka and Mishra, 2021) may form phase-separated droplets within the IC with the help of non-coding RNA sequences (Farabella et al., 2021; Razin and Gavrilov, 2021). Like assembled macromolecular machineries, such condensates are excluded by size from the INC and may serve as storage vesicles for the formation of functional machineries. For example, scaffold attachment factor A (SAF-A), was observed in the ANC (Smeets et al., 2014). Transitions between liquid and solid states of chromatin ensembles are still not well understood (Hansen et al., 2021). Whereas constrained local motions of nucleosomes occur on the seconds timescale, solid chromatin at mesoscale does not mix with its environment on the minutes to hours timescale. Phase-separation has been implicated in euchromatin and heterochromatin formation in a complex interplay with epigenetic mechanisms (Rowley and Corces, 2018). Chromosome territories apparently form a polymer network with material properties of an elastic hydrogel (Hansen et al., 2021). While a liquid state of 200 nm sized chromatin domains would enable the efficient activation of genes through diffusion-mediated, long-range regulatory contacts, the possibility that 200 nm domains exist as a solid gel state and exert gel-like sieving effects may also be considered. Evidence for a gel-like chromatin organization at mesoscale suggests a porous structure of the chromatin landscape with sieving effects affected by global or linear shapes of diffusing macromolecules (Cai et al., 2015; Pluen et al., 1999; Seelbinder et al., 2021). Phase-separated droplets, carrying an assembly of macromolecules (proteins, non-coding RNAs) required for specific nuclear functions, may interact with chromatin domains with gel-like properties. Such complex interactions are not explained by models of an exclusively size-dependent accessibility. For example, the penetration of small macromolecules assembled within such droplets into the interior of dense chromatin may be prevented, whereas percolation of individual, larger macromolecules is still possible.

A recent polymeric model of higher order chromatin arrangements at mesoscale was based on phase separation of chromatin, considering two key features, namely chromatin self-attraction and its binding to the peripheral lamina (Amiad-Pavlov et al., 2021; Bajpai et al., 2021). It offers a biophysical mechanism for the overall determination of nuclear chromatin density and hence accessibility by the regulation of lamina-chromatin interaction strength. Such a quantitative numerical model may help in part to explain physical properties of the ANC-INC model (Cremer et al., 2020b; Cremer et al., 2020a; Schmid et al., 2017). Other polymeric models provide similar interaction principles to control the density and hence accessibility of individual gene domains (Chiariello et al., 2016). Computer models of chromatin nanostructures performed under various biochemical assumptions, such as histone linker length, Mg^2+^ concentration, acetylation, histone interactions, or epigenetic marks argue for highly complex nanostructures containing relatively open parts as well as small regions with an apparent chromatin density sufficiently high to constrain the access of individual macromolecules, likely preventing the penetration of larger aggregates (Bascom et al., 2016; Odenheimer et al., 2009; Paulsen et al., 2018; Rosa and Zimmer, 2014). Schlick and colleagues (Bascom et al., 2019) performed numerical simulations based on Hi-C and epigenetic data of the 55 kbp-sized HOXC gene for a series of 3D configurations representing inactive and active chromatin states, compare Figure S3 in (Bascom et al., 2019). Extended model configurations show highly condensed clusters of densely packed nucleosomes connected by stretches of nucleosome-free DNA, i.e., highly accessible sites. From the volumes within the enveloping surfaces of these ensembles, one can estimate average densities ranging from ∼7 Mbp/µm^3^ to ∼20 Mbp/µm^3^. These average estimates, however, do not allow conclusions about the actual densities of structures below the resolution limit. Direct imaging of such structures remains an unsolved challenge and would likely require an intranuclear 3D structural resolution of a few nm.

Constrained spatial accessibility due to dense chromatin compaction of the INC is not the only barrier preventing/attenuating the diffusion of proteins and RNAs from the nucleosome free space of the ANC into the INC (Imai et al., 2017). Other features, such as electric charge (Cremer et al., 2000) and the enrichment of macromolecules within phase-separated droplets, may also affect the accessibility of chromatin ensembles. In addition, for more sophisticated models of chromatin accessibility, it would be important to achieve more information about the space-time distribution of nucleosomes (Baldi et al., 2020; Kharerin et al., 2020). To which extent dense convolutes of naked DNA may exist in the ANC has remained elusive.

True-to-scale DNA density maps should be complemented by similar density maps based on nucleosome positions as an essential basis for modeling of chromatin accessibility. Ideally, one would like to follow the dynamics of any target sequence of interest within a given chromatin ensemble and explore in living cells the dynamic compaction of individual loops or nanodomains comprising a few kb and the potential opening and closing of IC-channels. The generation of such space-time nuclear maps requires the development of time-resolved super-resolution microscopy approaches of live cells (Chu et al., 2021; Ricci et al., 2015; Shechtman et al., 2017; Wombacher et al., 2010; Xiang et al., 2018). The space-time visualization of chromatin ensembles is essential for studies to solve the question of intermingling loops between neighboring CTs (Branco and Pombo, 2006; Rouquette et al., 2010). Furthermore, an appropriate toolset should allow the integration of epigenetic marks, RNAs and non-histone proteins into individual chromatin ensembles across the entire nuclear landscape (Barth et al., 2020; Ma et al., 2016). Steadily improved, color-coded, high-resolution maps of nuclear landscapes are ideally suited to incorporate huge datasets of additional genetic and epigenetic information in an intelligible manner, providing direct insight into the interdependence of structural and functional changes (Gorkin et al., 2020; Lohoff et al., 2022; Yang et al., 2021). This is a long way to go not only with regard to the challenges of developing high resolution 4D live cell microscopy but also in view of the requirements of comprehensive, high-throughput 4D image analyses of large cell numbers. Changes of nuclear landscapes during differentiation and development of numerous cell types provide an important enrichment to the Human Biomolecular Atlas Program (Snyder et al., 2019). Evolutionary comparisons will shed new light on the origin of cell nuclei (Cremer et al., 2018) and the evolution of metazoan chromosomes (Simakov et al., 2022). Based on the emerging evidence that physical compaction and decompaction of chromatin nanodomains play a major role in gene regulation not only in eukaryotes but also in prokaryotes (Swygert et al., 2021), it is interesting to explore to which extent differences between high-resolution DNA density maps of different cell types (Yang et al., 2021) correlate with and may predict global transcriptional differences. Arguably, 3D nuclear arrangements of active and silent genes and their regulatory sequences within active and inactive nuclear compartments may be closely related with a cell’s transcriptome and protein interactome at a given time (Nussinov et al., 2021). If so, high resolution DNA-density maps of nuclei in a tissue may serve as structural markers reflecting different functional states (Lang et al., 2021).

## Limitations of the study

We currently lack equipment for an automated fBALM recording of large numbers of nuclei and algorithms for rapid image analyses. These prerequisites must be met for broad applications of our approach in research and diagnostic settings. The current structural resolution of SMLM (∼20 nm lateral, ∼100 nm axial) still prevents imaging of chromatin loops at the level of individual nucleosomes. Although our fixation protocols maintain 3D structures of chromatin ensembles to a remarkable extent down to the resolution limit of 3D-SIM (∼100 nm lateral, ∼250nm axial) (Markaki et al., 2013), we cannot rule out fixation artifacts at higher resolution. Live cell studies of nuclear landscapes with SMLM are currently impractical because the sequential recording of large numbers of frames for high resolution image reconstruction implies motion blur, while fewer frames imply sparse sampling and hence strongly reduced temporal and spatial resolution (Lelek et al., 2021; Schermelleh et al., 2019).

## Acknowledgments

Márton Gelléri, Shih-Ya Chen, Aleksander Szczurek, Jan Neumann, and Christoph Cremer were supported by the Boehringer Ingelheim Foundation and the Max Planck Institute for Chemistry, Mainz (Germany). Hilmar Strickfaden and Michael J. Hendzel were supported by grants from the Canadian Cancer Research Society (grant number CRISDI 2018 OG 23446) and the Canadian Institutes of Health Research (grant number PJT-148753). The Spinning Disk Confocal System (VisiScope, 5-Elements, IMB Microscopy Core Facility) and the High-Content Screening System (Opera Phenix, Perkin Elmer, IMB Microscopy Core Facility) were supported by the Deutsche Forschungsgemeinschaft (INST 247/912-1FUGG and INST 247/845-1 FUGG). We gratefully acknowledge the support of the Center of Advanced Light Microcopy (CALM) of the LMU Biocenter (headed by Heinrich Leonhardt and Hartmann Harz), for recording SIM images. We are obliged to many colleagues for helpful suggestions. In particular, we thank in alphabetical order Udo Birk, Giacomo Cavalli, Jonas Cremer, Mario Nicodemi, Thoru Pederson, Katharina Ribbeck, Karsten Rippe, Vassilis Roukos, Samuel Safran, and Nadine Vastenhouw for critical reading and discussion. We thank Sandra Ritz and all the members of the IMB Microscopy Core Facility for technical support.

## Author contributions

CC, HS, MG, MC and TC planned this study and wrote the manuscript. All authors contributed to the final text. S-Y C and CC developed the SMLM instrumentation; AS developed the fBALM approach and contributed experimental SMLM results. SMLM-images of nuclear landscapes were recorded by S-Y C, MG, HS, and AS, electron spectroscopic images by HS. Quantitative Voronoi-analyses of SMLM sections were carried out by MG, analysis of ESI images by HS and MH. Evaluation algorithms were provided by MG, JN, OK, FS, HS and JI. These authors also contributed to the various approaches of SMLM image displays and analyses. Microinjection experiments with fluorescent nanobeads and quantitative analyses were conducted by BH and HS. EdU-replication-labeling experiments with SMLM were performed by AS. MS carried out 3D SIM studies of EdU-pulse-chase experiments with guidance from YM.

## Material & Methods

### Cell cultures, fixation, and staining protocols

h-TERT BJ1 cells, an immortalized diploid human cell line from human foreskin fibroblasts (ATCC #CRL-2522); HeLa cells, a human cervix carcinoma derived cell line with a near-triploid karyotype (Landry et al., 2013), and C3H 10T1/2 cells, derived from a murine embryo fibroblast cell line (ATCC #CCL-226) (Bairstow and Heidelberger, 1975) were cultured in DMEM with 5% CO_2_ at 37 °C. Undifferentiated female mouse embryonic stem cells (ESC line 16.7, 42,XX; +6;+8, kindly provided by J. Lee, Harvard Medical School) were cultivated under feeder-free conditions on gelatinized cover slips in KO-DMEM (Invitrogen) as described in details in (Smeets et al., 2014). After fixation with formaldehyde (4%) for 15-30 min, cells were kept in Dulbecco’s Phosphate Buffered Saline (DPBS, Sigma Aldrich) and incubated with 0.5 U/ml RNase A and 20 U/ml RNase T1 (Ambion, USA) for 1 h at 37 °C. It is essential to keep living cells in their normal osmotic environment (Albiez et al., 2006) and avoid drying out of preparations during fixation and all subsequent steps until embedding for microscopic examination in order to maintain the shape, volume and internal structure of cells as best as possible. We employed protocols that maintain 3D nuclear shape and topography of CTs, nuclear speckles, replication domains and 3D positions of genes to a remarkable extent down to the lateral and axial resolution limit of 3D-SIM (x,y ∼ 100 nm and z ∼ 250nm) (Markaki et al., 2013). A recent study performed with live and fixed Drosophila larva muscle nuclei found that the live nucleus preserved its volume and ellipsoidal shape, whereas the fixed nucleus exhibited substantial flattening, forming a disk-like shape (Amiad-Pavlov et al., 2021). This flattening effect, however was not observed in our present study (Supplemental Figure S8). In 3D-SIM studies live and fixed cells showed a high degree of similarity with regard to chromatin clusters, sites of decondensed chromatin, IC lacunae and IC channels leading to nuclear pores. A significant impact of fixation artifacts on large-scale chromatin organization was excluded in 3D-SIM live-cell experiments with HeLa cells stably expressing histone H2B-GFP (see additional file 8 in (Smeets et al., 2014). Still, we cannot exclude the possibility of fixation artifacts of active and inactive chromatin ensembles at nanoscale below the resolution limit of 3D SIM.

For DNA staining the following dyes were used: Sytox Orange or Vybrant Violet for SMLM-fBALM (fluctuation assisted Binding Associated Localization Microscopy Mode (Chen et al., 2018; Szczurek et al., 2018; Szczurek et al., 2017; Zurek-Biesiada et al., 2016), see below. These dyes were present at a concentration of 0,5 µM throughout the whole imaging procedure. For other microscopic approaches (wide-field, confocal and structure illumination), nuclei were pre-stained with DAPI (5 µg/ml for at least 5 min) and Hoechst (10µg/ml; for at least 5 min) and washed in PBS before use. With a size in the 1 nm range, the accessibility of all dyes should not be substantially impaired by chromatin compaction per se.

### Determination of cell cycle & nuclear morphology

Determination of cell cycle stage and nuclear morphology of BJ1 and HeLa cells was performed using a spinning disk Opera Phenix Plus High-Content Screening System (PerkinElmer), according to (Roukos et al., 2015), with Sytox Orange as a DNA stain. If not stated otherwise, nuclei corresponding to G1 were used for SMLM-fBALM imaging (Supplemental Figure S2). Imaging was performed in wide field mode with a 20x, 0.4 NA objective to capture fluorescence from interphase cells and mitotic cells without the need of recording a z-stack. In each well, multiple images were acquired, covering approximately 80 percent of a single well resulting in 20,000 to 80,000 imaged cells per well. Nuclear area and fluorescence intensity of Sytox Orange was analyzed by segmentation with Harmony software (PerkinElmer). Nuclei that were touching the image border were excluded from further analysis. The frequency distribution of Sytox Orange intensities showed discrete G1 and G2 cell cycle populations that allowed the classification of each cell nucleus into these classes.

### Microinjection of fluorescent nanobeads (FluoSpheres)

Microinjection of carboxylate-modified FluoSpheres (ThermoFisher Scientific) with a diameter of 20 nm and 40 nm into nuclei was performed on an Axiovert 200 M widefield microscope (Zeiss) with an injection system from Eppendorf. Needles were pulled from borosilicate glass capillaries with an outer diameter of 1.0 mm and an inner diameter of 0.5 mm using a micropipette puller (Sutter Instrument P-97; for details see (Hübner, 2020).

### Pulse-labeling of replication domains (RDs) with EdU

For SMLM experiments, RDs were identified following pulse-labeling of HeLa cells for 15 min with 5-Ethynyl-dU (EdU) at a final concentration of 10 μM as described (Cremer et al., 2020b; Kirmes et al., 2015). After fixation of cells with 4% formaldehyde/PBS for ∼20 min and permeabilization with 0.5% Triton X-100/PBS/0.02% Tween for 10 min, EdU was conjugated to azid modified AlexaFluor 555 and detected by click-chemistry according to manufactures instructions (https://www.thermofisher.com/). DNA was stained with Vybrant Violet (0.5 µM). For 3D-SIM experiments, RDs in mouse embryonic stem cells were identified by an EdU pulse of 10 min. Cells were fixed immediately thereafter with 2% formaldehyde, or after a chase of 20 - 80 minutes and permeabilized with 0.5% Triton X-100/PBS/0.02% Tween for 10 min. EdU was conjugated to azid modified AlexaFluor 488 and detected by click-chemistry according to manufactures instructions (https://www.thermofisher.com/). After post-fixation in 4% formaldehyde for 10 min DNA was counterstained with DAPI (5 µg/ml).

### Distances between surfaces of uniformly arranged nucleosomes in 3D space estimated for different DNA densities

We assumed that the entire genome was fully packaged with nucleosomes, i.e. that each consecutive 200 bp DNA segment carried a nucleosome with ∼150 bp of DNA, wrapped around the core histone complex, and a linker length of ∼50 bp between any two linearly adjacent nucleosomes located on the same ∼10 nm nucleosomal chain. In case of fully extended linker DNA, this means a maximum surface-to-surface distance of ∼15 nm. Next, we determined the maximum number and spatial distribution of nucleosomes that can be placed at a given DNA density into a 1 µm^3^ cube under the highly simplified assumption of homogenously distributed, equidistant nucleosomes. Placing each nucleosome (represented by spheres with 10 nm diameter) with its center into a sub-cube, we calculated the center-to-center 3D distances between adjacent nucleosomes placed in the midst of these sub-cubes; from this, the minimum average 3D distances between the surfaces of spatially adjacent spherical nucleosomes were determined for average DNA densities of 5, 30, 60 and 150 Mbp/µm^3^. For example, a cube of 1 µm^3^ with an average DNA density of 150 Mbp/µm^3^ contains ∼750,000 nucleosomes placed at a mean 3D distance of ∼1 nm between the surfaces of any two nucleosome neighbors adjacent in 3D space. In contrast, a 1 µm^3^ cube with an average DNA density of 5 Mbp/µm^3^ contains only ∼25,000 nucleosomes with a minimum 3D distance of ∼24 nm. Note that many next nucleosome neighbors in 3D space may be located at different DNA sites of the same nucleosomal chain or on different spatially adjacent loops. This explains why the minimum average distance calculated for 5 Mbp/µm^3^ exceeds the maximum length of ∼15 nm for a fully stretched 50 bp DNA linker.

These estimates cannot replace sophisticated modeling, taking into consideration the constraints of polymer chain folding of 10 nm thick nucleosomal chains (Amitai and Holcman, 2017). Notably, highly compacted chromatin does not provide much freedom for local changes of accessibility resulting from a dynamic, non-uniform distribution of nucleosomes. Whereas the accessibility of the interior of a loop forming a dense nucleosome cluster is highly constrained (Maeshima et al., 2015), such constraints may barely matter in case of an open loop configuration (compare Figure 1c).

### Single Molecule Localization Microscopy (SMLM)

For studies of high-resolution DNA density maps with SMLM-fBALM (fluctuation-assisted Binding Activated localization microscopy) mode (Szczurek et al., 2018; Szczurek et al., 2017) an inverse custom-made localization microscope was equipped with three lasers emitting at 488 nm, 561 nm and 647 nm. The three lasers were combined with dichromatic reflectors. The laser beams were expanded by a pair of lenses and focused to the back focal plane of the objective lens (HCX PL APO 100x/NA 1.47 oil, Leica, Germany) by a tube lens. A quadband dichroic mirror was used to reflect the laser and to transmit emitted fluorescence light from the sample. In this setup, a Gaussian intensity distribution with a full width at half maximum (FWHM) of 50 µm was created in the focal plane of the objective lens.

fBALM relies on the spatial switching of fluorescent on/off states by adjusting the local environment of the DNA and DNA-binding sites. The chemical environment, in particular the pH value, allows control over the local, reversible DNA melting of double stranded into single stranded DNA. Sytox Orange binds to dsDNA, falls off after local melting from ssDNA, and rebinds to dsDNA, usually at another site. Unbound Sytox Orange molecules have a fluorescence up to several orders of magnitude lower than the emission of blinking DNA-bound molecules. Furthermore, unbound Sytox Orange molecules have a much larger mobility. Hence their weak fluorescence emission forms a rather homogeneous background which can be removed by appropriate image analysis tools (Kaufmann et al., 2009). In summary, fBALM provides both a substantial reduction of background combined with a very high number of blinking events of dsDNA bound fluorophores, which is only limited as a consequence of either complete bleaching of the entire pool of Sytox Orange molecules, a change of the chemical environment, or of an optical instability of the SMLM setup.

SMLM-fBALM allowed the determination of single-molecule positions (SMPs) of Sytox Orange molecules in nuclei with a typical average precision of σ_xy_ ∼8 - 15 nm (object plane; best values around 2 nm in heterochromatic regions), and an axial precision σ_z_ down to about 50 nm. From these figures, an average lateral optical resolution OR_xy_ (smallest lateral distance detectable between two molecule positions) of 20 nm was obtained for σ_xy_ = 8.5 nm, and around 100 nm for OR_z_ along the optical axis. For molecule positions with a localization precision (object plane, xy) of 2 nm, the best OR_xy_ value estimated was around 5 nm. Experimentally, in nuclear sections of HeLa nuclei, selected single molecule positions with σ_xy_ ≤5 nm and mutual xy-distances of 11 nm and of 14 nm, respectively, were clearly distinguished. For the object plane coordinates with σ_xy_ = 8.5 nm and d (SMPs/nuclear area = 10^7^/100 µm^2^ = 0.1/nm^2^), the average structural resolution R (as a measure for the smallest observable structure) was estimated according to R = [(2.35 σ_xy_)^2^ + (4/d)]^0.5^ (Cremer et al., 2017). From this, R = 21 nm was obtained if all 10 million molecule positions localized are included in a 600 nm optical section. For a thinner optical section of about 100 nm with 1.5 x10^6^ molecule positions, R = 26 nm was estimated. If in addition the average drift (after correction) is included (Zurek-Biesiada et al., 2016), then the average R increases from 21 nm to 23 nm, and from 26 nm to 28 nm, respectively.

Selected HeLa and BJ1 cells were placed in the center of the field of view of the inverted SMLM-microscope. Nuclei were exposed to the 561 nm laser at an estimated illumination intensity of about 1 kW/cm^2^. Fluorescence was collected by the objective lens and was separated from the excitation laser by a dichromatic beam splitter and appropriate emission filters. The emission light was focused by a second tube lens and imaged on an sCMOS camera (PCO edge 4.2) with a pixel size corresponding to 65 nm in the object plane, setting the exposure time to 50 ms/frame. Up to 50,000 frames with a total of ∼10 million blinking events were recorded in a single ∼600 nm thick light optical section and localized with appropriate localization algorithms (Cremer et al., 2017; Kaufmann et al., 2009). The possibility that the same fluorophore was recorded twice (double blinking event) was considered in cases where two signals recorded in subsequent image frames had a distance smaller than 20 nm. These rare events were excluded from further image analysis as described below. To localize SMPs from a thinner section than 600 nm, achieved with conventional wide-field resolution, a cylindrical lens was placed in front of the sCMOS camera (Chen et al., 2018) to induce an astigmatism (Huang et al., 2008). In this way it was possible to assign SMPs to positions ± 50 nm above or below the focal plane and thus filter out the SMPs in a ca. 100 nm thick optical section (assumed to be ca. 1/6th of the total number of SMPs in the 600 nm optical section).

For recording of DNA densities and replication domains (RDs) with dual-color SMLM, a custom-built, upright-setup (Reymann et al., 2008) was used, equipped with lasers for excitation wavelengths 405 nm and 491 nm, respectively (Zurek-Biesiada et al., 2016). EdU-labeled RDs were conjugated to AlexaFluor 555 and recorded first (5,000 SMLM frames). Vybrants DyeCycle™ Violet (VdcV) was used to label the non-replicated DNA (Zurek-Biesiada et al., 2016). After photoconversion with 405 nm, Vybrant Violet was excited to ‘blink‘ by intense 491 nm illumination, and the emitted green fluorescence was registered (30,000 frames).

### Estimation of true-to-scale DNA density values

The procedure we have used to achieve true-to-scale DNA density values is illustrated in Supplemental Figure S3 using the BJ1 light optical section presented in Figure 2. The reliability of 3D DNA density values for a given chromatin ensemble critically depends on the determination of both its absolute DNA content and its volume. Several essential requirements for precise measurements of these values need to be considered.

First, our concept assumes that Sytox Orange (Szczurek et al., 2018) binds randomly, i.e. without base-pair preference, to the nuclear dsDNA, and can be localized with high precision by SMLM. The lateral nuclear positions of SMPs (within the object plane) were determined with an average precision of 8 - 15 nm. This requires extreme mechanical stability of the microscopic setup (Chen, 2020; Chen et al., 2019). The determination of the axial positions of SMPs (Chen et al., 2018) was not further defined, but assumed to reflect the thickness of the light optical section, which was about 600 nm in the example HeLa cell nucleus (Figure 3). In the case of the BJ1 nucleus (Figure 2) and HeLa nucleus (Figure 3), the use of an astigmatic lens (Huang et al., 2008) resulted in an improved axial localization precision of SMPs down to about 50 nm in a 100 nm thick optical section. Under optimal conditions, lateral nuclear SMP distance differences <5 nm were detected. The pixel size for the density determination was adapted to reflect the mean localization precision of the SMLM measurements. For volume estimates of light optical sections, we assumed a thickness of 100 nm in SMLM setups with an achromatic lens and of 600 nm without such a lens.

Second, for estimates of the total DNA content of an entire nucleus or a given light optical section, it is necessary to determine the interphase stage of the studied cell. Cell cycle measurements (see above) indicated that the cells subjected to fBALM were in G1. In the case of cells with an abnormal chromosome content, such as HeLa cancer cells, it is preferable to measure the DNA content of a single, studied cell nucleus (not performed here) or to provide an estimate based on knowledge of the karyotype, though this karyotype may differ from cell to cell. For BJ1 cells we assumed a diploid karyotype (2n), and for HeLa cells a near triploid karyotype (3n).

Third, the calibration of absolute DNA density values should ideally be based on the total number of SMPs mapped within an entire cell nucleus. This total number then reflects the total DNA content. Accordingly, the fraction of genomic DNA represented by a single SMP can be calculated. Ideally, this approach would require high resolution, optical serial sectioning of a given nucleus with SMLM-fBALM in order to sum up the total number of blinking events and determine its shape and volume together with its total DNA content. In practice, the decrease of blinking efficiency due to a deterioration of the chemical switching conditions (Szczurek et al., 2017) during the long registration times prevented serial SMLM sections of entire nuclei with high structural resolution. For this reason, we typically determined the height of the nucleus given by the distance between the top and bottom section and performed a single SMLM midsection from nuclei with a high average SMP density. For simplicity, we assumed a homogeneous DNA distribution, yielding a linear relationship between volume and DNA content. Furthermore, we assumed that the shape of a nucleus conforms to a cylinder with a circular or elliptical base and the specified height.

Fourth, calibration of the amount of DNA represented by each SMP in a given section allows estimates of the absolute DNA density of a given chromatin ensemble, recorded in this section. For this purpose, it is necessary to count the number of SMPs attributed to such an ensemble. This is easy in cases, where the outline of an ensemble can be precisely determined, for example after FISH of DNA targets with specific probes (Bintu et al., 2018; Weiland et al., 2011). In the absence of further information, however, demarcation of individual ensembles, say a chromatin loop, a chromatin domain, or a chromosome territory, from neighboring structures, is difficult and often impossible. Using a customized version of a published Voronoi algorithm (Andronov et al., 2016), we dissected the cloud of SMPs into Voronoi areas limited by lines with equal distances to neighboring SMPs. The smaller the distances between neighboring SMPs, the smaller were the defined Voronoi areas, and the higher was the estimated density of SMPs. This approach was performed without any density thresholding based on biological assumptions about chromatin ensembles with specific structural or functional properties. Based on Voronoi area frequencies we generated a histogram of the DNA density distribution across a given optical section. Based on the calibration between the number of SMPs and the corresponding DNA content, histograms of Voronoi area frequencies were converted into a color-coded map of absolute DNA densities (Mbp/µm^3^). However, this advantage was achieved at a cost: Voronoi tessellation is particularly useful to detect local DNA densities, likely corresponding to small nucleosome clusters, but fails to detect chromatin loops formed by extended ∼10 nm thick nucleosomal chains or naked DNA.

### 3D-Structured Illumination Microscopy (SIM)

Super-resolution imaging of fixed cell samples with 3D-SIM was performed on a DeltaVision OMX V3 system (Applied Precision Imaging/GE Healthcare) equipped with a 100x/1.40 NA PlanApo oil immersion objective (Olympus), Cascade II:512 EMCCD cameras (Photometrics) with 405 and 488 nm diode lasers. 3D-SIM image stacks were acquired with a z-distance of 125 nm and with 15 raw images per plane (5 phases, 3 angles). The raw data was then computationally reconstructed using Wiener filter settings 0.002 and channel-specifically measured optical transfer functions (OTFs) using the softWoRX 6.0 software package (Applied Precision) to obtain a super-resolution 3D image stack with a lateral (xy) resolution of ∼120 nm and an axial (z) resolution of ∼300 nm. The level of spherical aberration was minimized and matched to the respective OTFs using immersion oil with different refractive indices (RI)(Smeets et al., 2014). Images from the different color channels were registered with alignment parameters obtained from calibration measurements with 0.2 µm diameter TetraSpeck beads (Invitrogen). The reconstruction process generates 32-bit data sets with the pixel number doubled in the lateral axes, and the lateral pixel size halved from 80 nm to 40 nm in order to meet the Nyquist sampling criterion. To normalize all image stacks for subsequent image processing and data analysis, the stack-specific mode gray value (representing the peak of the background noise) was subtracted, negative values discarded and finally the format converted to 16-bit composite tiff-stacks using a custom-made script based on ImageJ (http://rsbweb.nih.gov/ij).

### Confocal microscopy of microinjected nanobeads

For confocal microscopy of microinjected nanobeads, a laser scanning and a spinning disk confocal microscope were used. The TCS SP5 DMI 6000 CS CLSM microscope (Leica Microsystems, Mannheim/Germany) was equipped with a HCX Plan Apochromat Lambda Blue 63x/1.4 oil objective lens and various laser sources, the excitation wavelengths ranging from 405 nm to 633 nm. Spinning disk confocal microscopy was performed on an Ultraview system (PerkinElmer) equipped with 405 nm, 488 nm, and 561 nm diode lasers using a 100x/1.4 oil Apochromat (Zeiss) objective lens.

### Electron Spectroscopic Imaging (ESI)

C3H 10T1/2 growing in 35mm MatTek glass bottom dishes cells were fixed in 0.1 M Phosphate Buffer containing 4% paraformaldehyde and 2% glutaraldehyde for 30 min. Then the cells were permeabilized with 0.1% Triton X-100 for 15 min. After washing with 1xPBS the cells were treated with RNAse as described above. After a dehydration series (50%, 70%, 90%,100%) each 30 min, the cells were infiltrated with acrylic resin (LR White) (1:1 mixture with 100% ethanol for 4h and in pure LR White overnight). Placing an LR White filled 2 ml Eppendorf tube on top of the cells and polymerizing the acrylic resin for 48h at 65°C in the oven, blocks were created that could after detaching from the glass be trimmed and cut with a Diatome diamond knife on a Leica Ultracut UC6 microtome (Strickfaden et al., 2015). 100 nm ultrathin sections were picked up on 300 Mesh copper grids and carbon coated. Cells were located in a JEOL F2100 transmission electron microscope. Phosphorous and nitrogen elemental maps were recorded using a GATAN Quantum post column electron energy loss spectrometer equipped with a GATAN UltraSCAN 1000 CCD camera. For phosphorous elemental maps, post-edge pictures were recorded at 162.5 eV (slit width 20 eV) and pre-edge images at 119.5 eV and 99.5 eV (slit width 20 eV each) (3-window method). For nitrogen elemental maps, post-edge pictures were recorded at 416 eV (slit width) and pre-edge pictures were recorded at 416 eV and 383 eV (slit width 30 eV each). Phosphorous and nitrogen elemental maps were computed in Digital Micrograph by GATAN.

## Supplementary Information

**Supplemental Figure S1.**
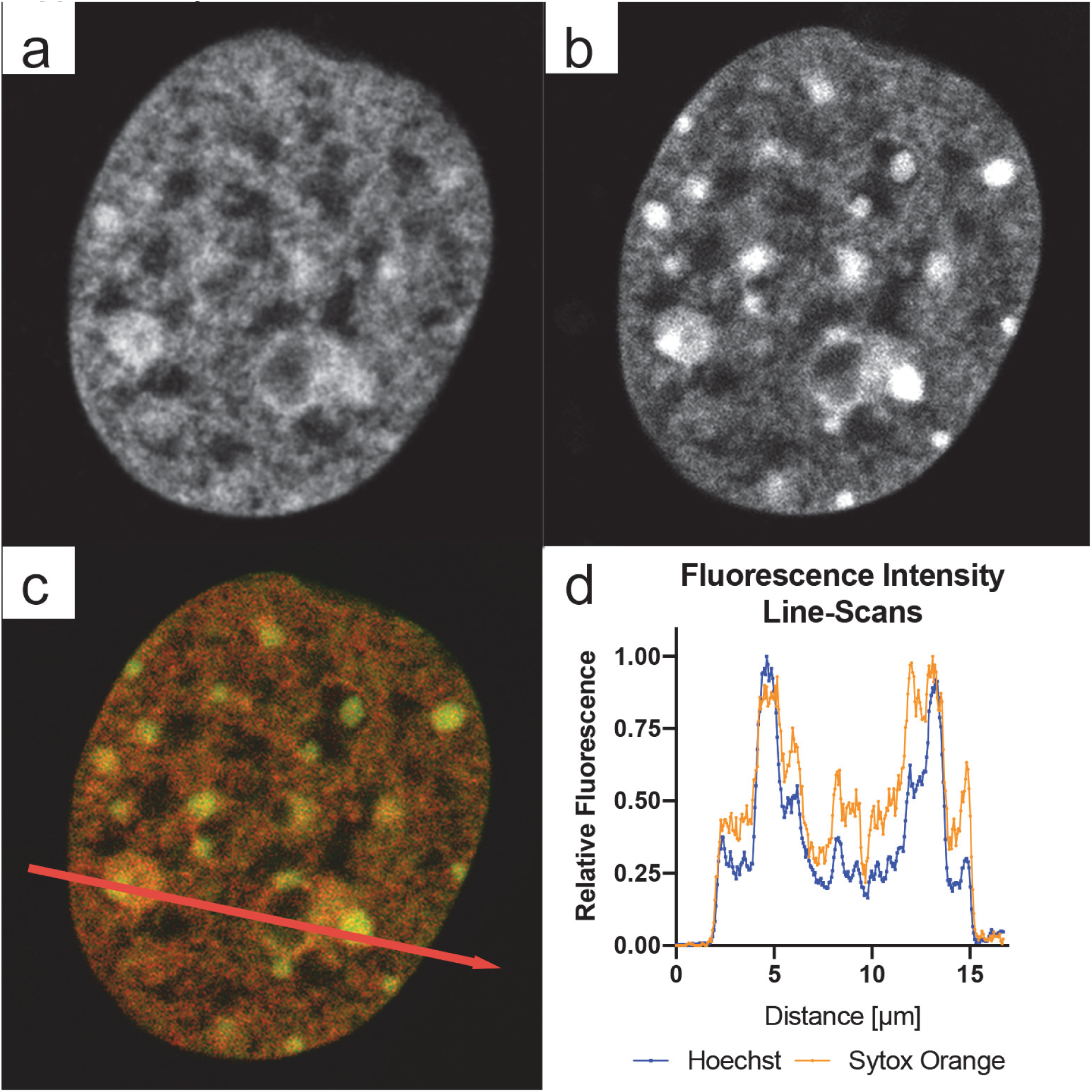
Comparison of Sytox Orange and Hoechst 33342 DNA counterstains. Confocal light-optical section of a C3H10T1/2 nucleus after RNase digestion and double-staining with the DNA dyes: **a** Sytox Orange, **b** Hoechst 33342, **c** merged image. **d** Fluorescence intensity distribution of both signals along the line through the nucleus shown ina-c. Note the improved staining of Sytox Orange in euchromatic regions.

**Supplemental Figure S2.**
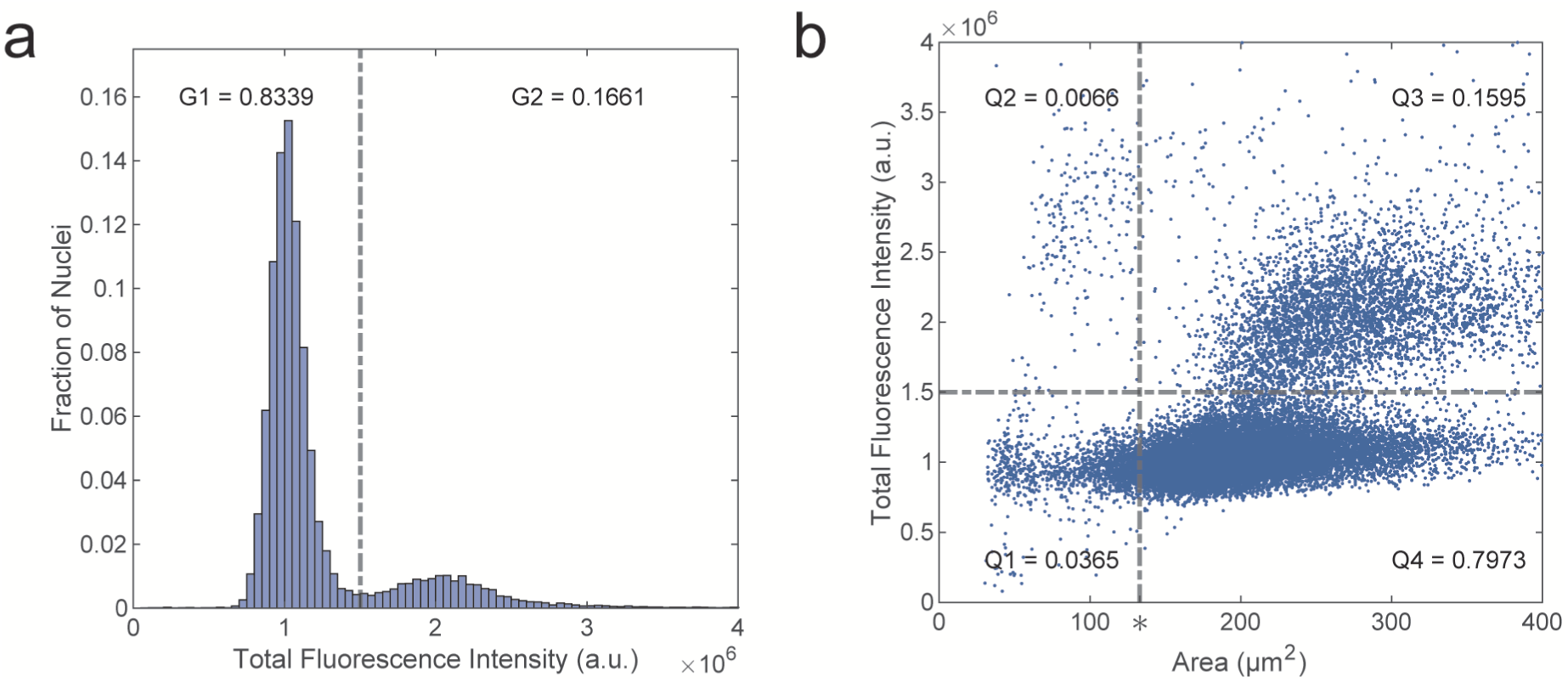
Cell cycle estimation of BJ1 nuclei. **a** Histogram of total fluorescence intensities of BJ1 nuclei stained with Sytox Orange. For an estimate of the cell cycle stage, cell nuclei with a total fluorescence intensity below or equal to 1.5 x 10^6^ a.u. (arbitrary units) were attributed to G1, and nuclei above this threshold were attributed to G2. According to this, 83 percent of the cells were found in G1 and 17 % in G2. **b** Total fluorescence intensity plotted vs nuclear area. The horizontal dashed line denotes the total fluorescence intensity that was used to discriminate G1 and G2 in (a). The vertical dashed line indicates the size of the BJ1 cell shown in Fig. 2. Q1 to Q4 give the fraction of nuclei found in the 4 quadrants.

**Supplemental Figure S3.**
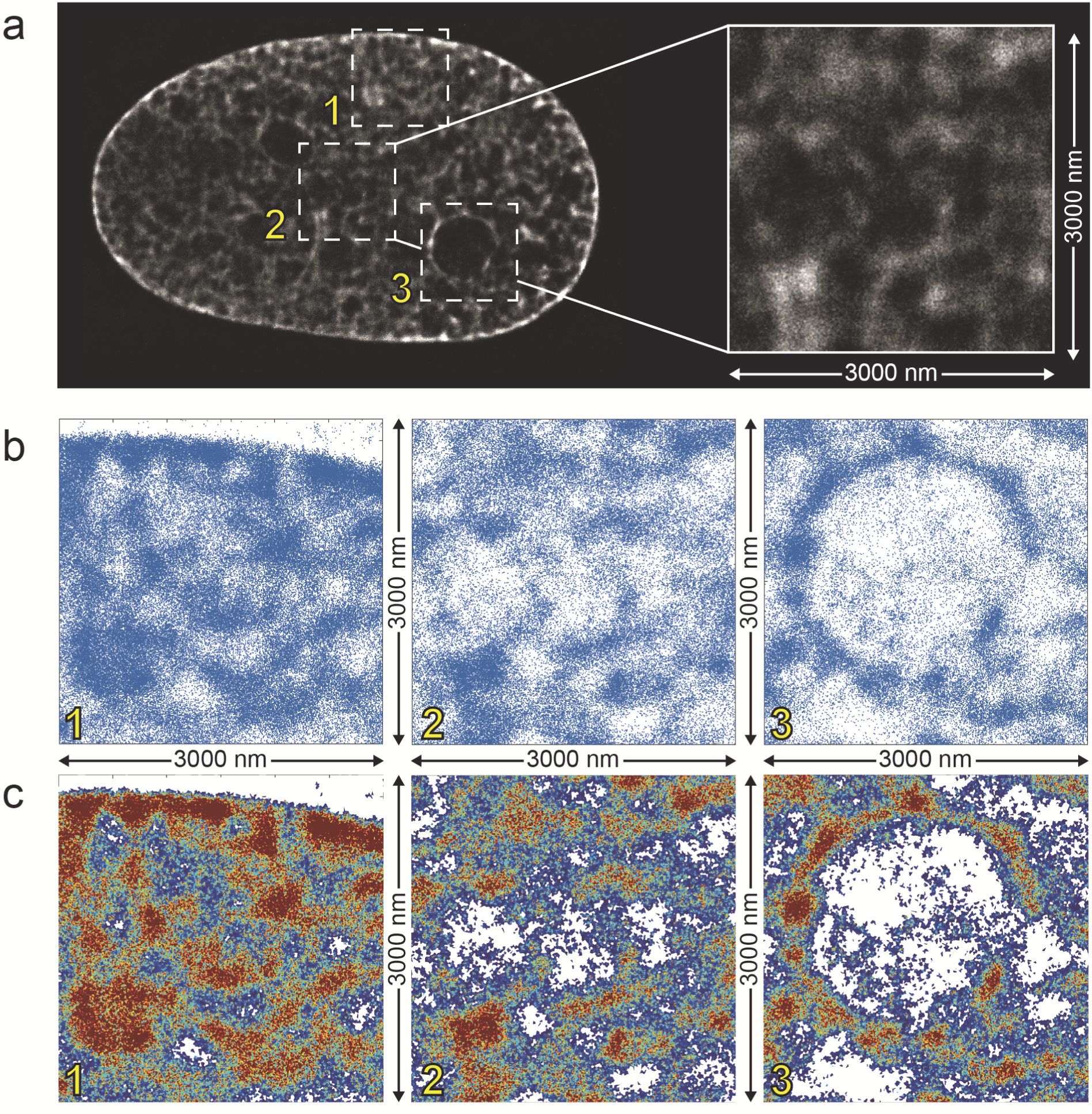
Processing of SMLM images and generation of high-resolution nuclear landscapes. **a** SMLM-fBALM image of a mid-plane optical section through a BJ1 cell nucleus stained with Sytox Orange yielding about 10 million SMPs with a lateral structural resolution of about 20 nm and an axial resolution of about 600 nm. SMPs were binned in pixels with 10 nm edge length. For this overview, a region was chosen with densely arranged SMPs corresponding to the INC. The three boxes mark regions at the nuclear periphery (1), in the nuclear center (2) and a nucleolus (3). Supplemental Video S1a shows a movie of the enlarged box 2 in (a) with SMPs at increasing density and structural resolution; Supplemental Video S1b provides a view on the rotating 600 nm thick section. **b, 1-3** Enlargements of boxes 1-3 represent 2D localization maps of all individual SMPs in 100 nm thick optical subsections with an average lateral localization precision of ca. 8 nm and ca. 50 nm along the optical axis (see Results for explanation, how 100 nm thick subsections were achieved from the 600 nm thick primary optical section). **c, 1-3** The same boxes as above are presented here after Voronoi-tessellation as true-to-scale color-coded DNA density maps (Mbp/µm^3^); compare DNA density scale bar in Figure 2, where the identical panel is shown in Fig. 2b1-3. For calculations of 3D DNA densities see Results.

**Supplemental Figure S4.**
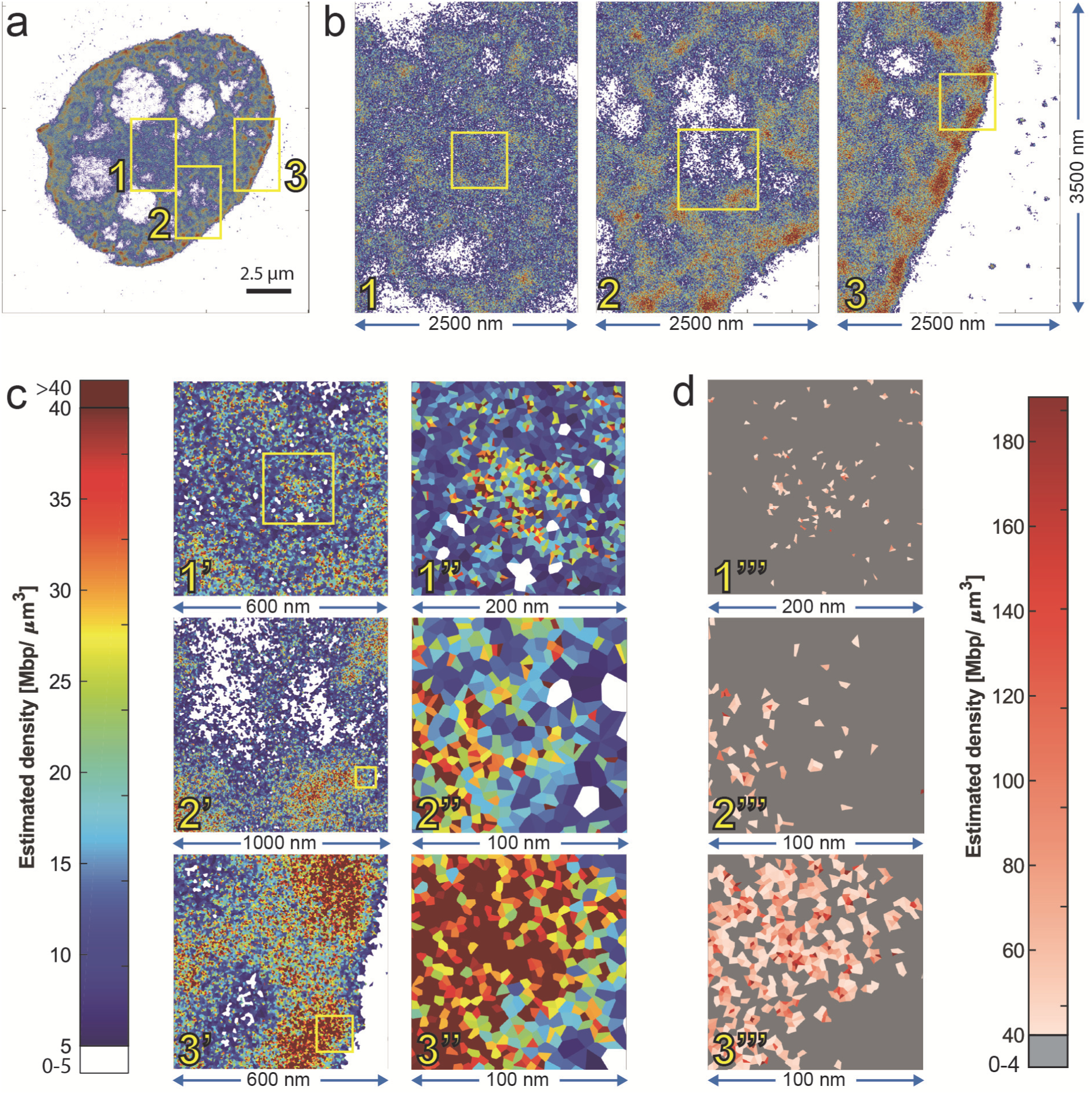
High-resolution, true-to-scale DNA density landscape of another HeLa cell nucleus. **a** Voronoi-tessellated SMLM-fBALM image from a mid-optical section through another HeLa cell nucleus shown for a phenotypic comparison with the nucleus in Fig.3. b, 1-3 Magnifications of the three boxed areas in (a); edge lengths 2.5 µm x 3.5 µm. **c** color map, indicating 0-5 Mbp/µm^3^ (white), and the range from 5 to (dark blue) to >40 Mbp/µm^3^ (dark red) (as described in Figure 2). **c, 1’-3’** Further magnification of boxed areas in (b,1-3); edge length 600 nm(1’), 1 µm (2’), and 600 nm (3’). **c, 1’’-3’’** Further magnification of boxed areas in c,1’- 3’; edge length 200 nm (1’’) and 100 nm in (2’’) and (3’’). **d, 1’’’- 3’’’** presents the same regions as in c,1’’- 3’’ using another color code (also described in Figure 2), demonstrating the range of apparent DNA densities above 40 Mbp/µm^3^ up (light red) up to maximum densities of ∼180 Mbp/µm^3^ (dark red), whereas all Voronoi areas assigned to densities between 0 and 40 Mbp/µm^3^ are grey colored. Note: Image recording of this nucleus was performed without an astigmatic lens, yielding a thickness of the optical section of about 600 nm. Within this optical section, about 4 million blinking DNA-bound Sytox Orange molecules were registered in 25,000 consecutive frames. The midsection of the nucleus presented in Figure 3 is smaller (88 µm^2^) than the midsection area of this nucleus (96 µm^2^), which arguably may reflect a DNA density landscape of a transcriptionally more active cell nucleus.

**Supplemental Figure S5.**
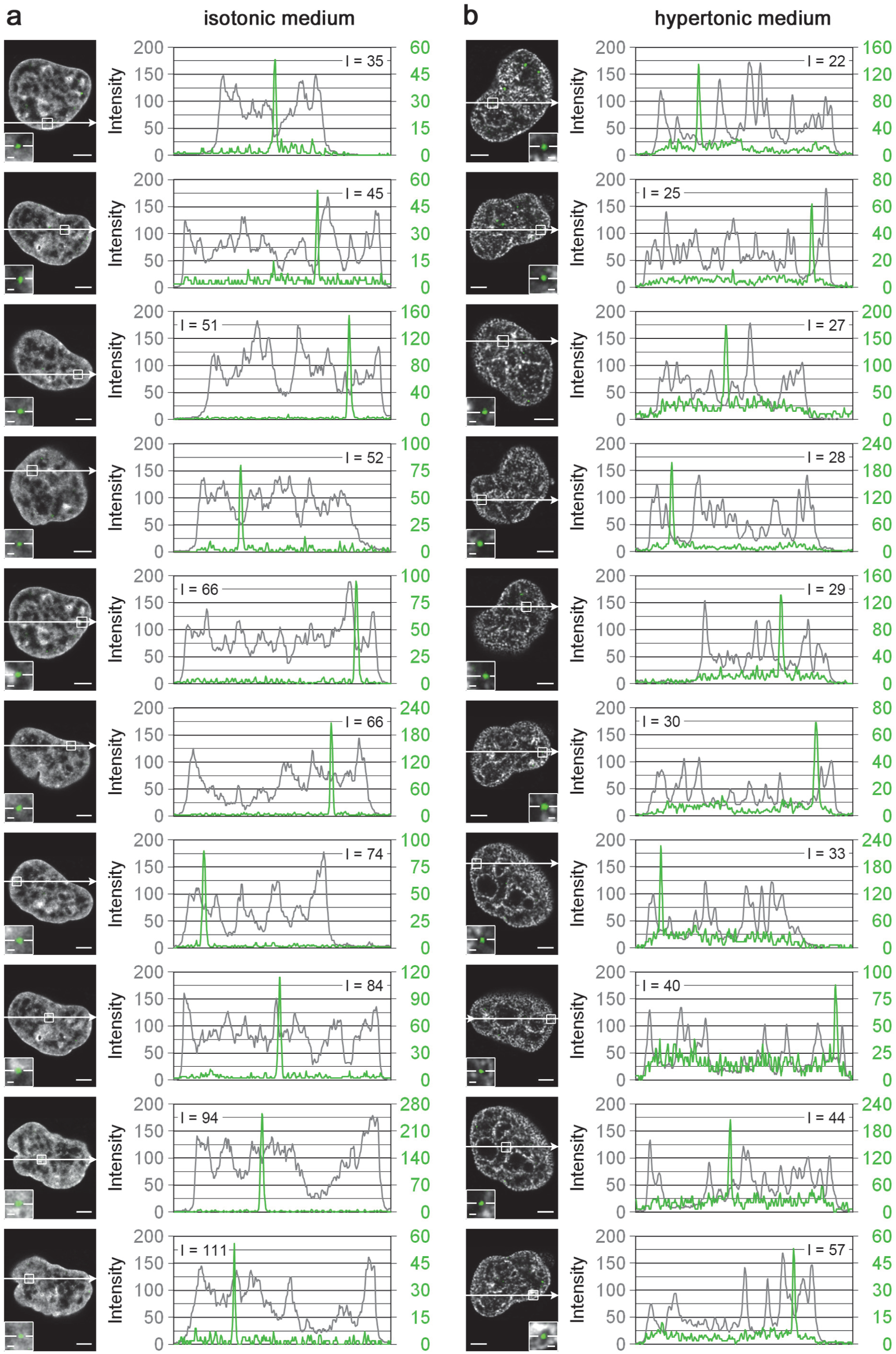
Localization of microinjected 20 nm nanobeads. Localization of nanobeads (with 20 nm diameter) in nuclei of HeLa cells kept in **(a)** isotonic medium with normally condensed chromatin (NCC) or **(b)** hypertonic medium with hyper-condensed chromatin (HCC). The confocal sections (left columns in a and b) show the position of the intensity line plotted at the right; the arrow denotes the scanning direction. The area of the bead is magnified in the inset. Scale bars: 4 μm, 0.5 μm in insets. For each nucleus, line scans were made through a given bead (green line) and through the normalized DAPI intensity image (black line). The relative DNA density (I; grey values 0 - 255) present at the site of a nanobead was determined in the confocal section with the brightest spot. Nanobeads are localized within valleys of the DNA density landscape and sometimes at the slope of DNA density peaks.

**Supplemental Figure S6.**
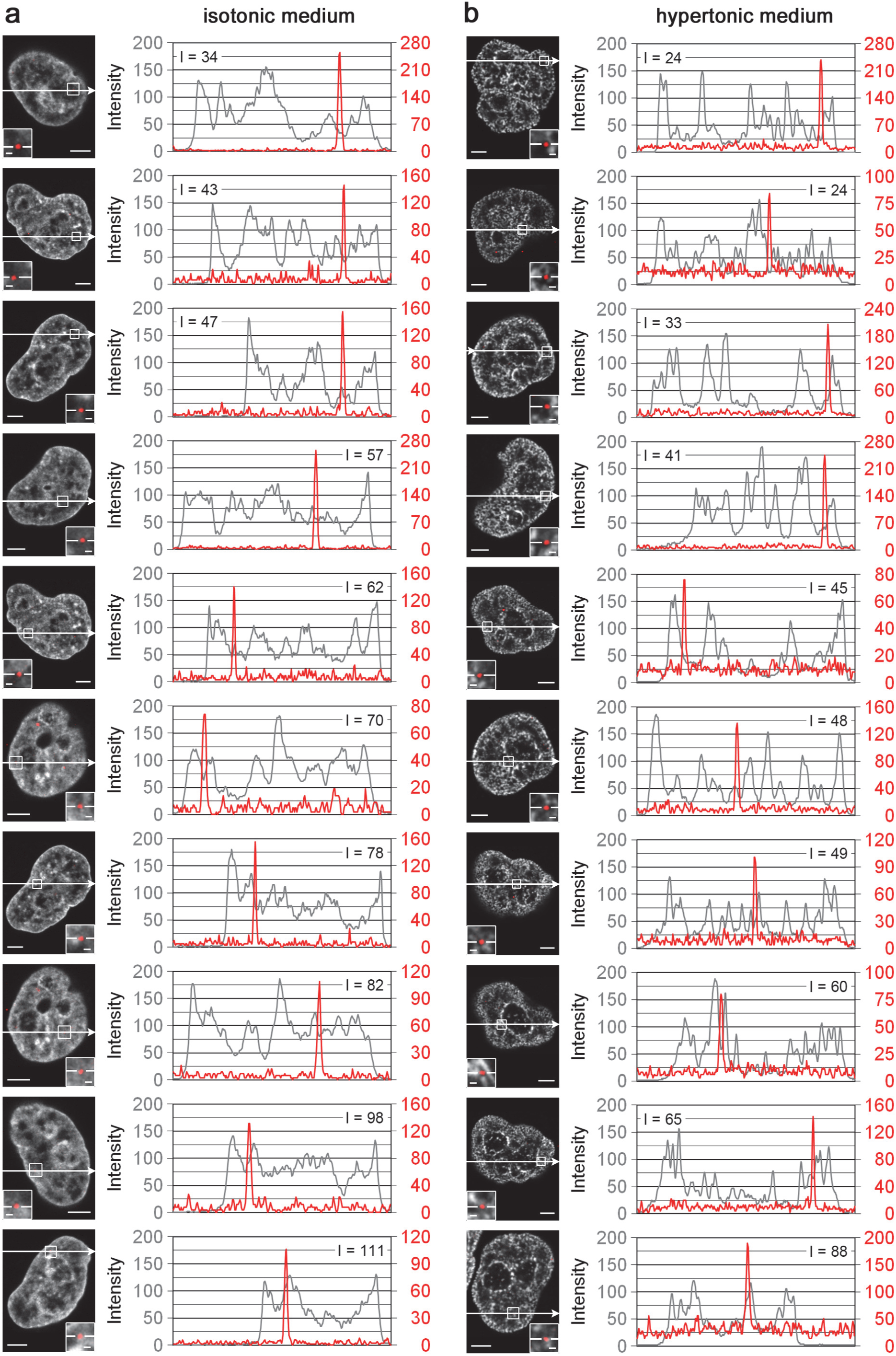
Localization of microinjected 40 nm nanobeads. Localization of nanobeads (with 40 nm diameter) in nuclei of HeLa cells kept **(a)** in isotonic medium with normally condensed chromatin (NCC) or **(b)** hypertonic medium with hyper-condensed chromatin (HCC). Nanobeads are localized within valleys of the DNA density landscape and sometimes at the slope of DNA density peaks. For further details see legend to Supplemental Figure S5.

**Supplemental Figure S7.**
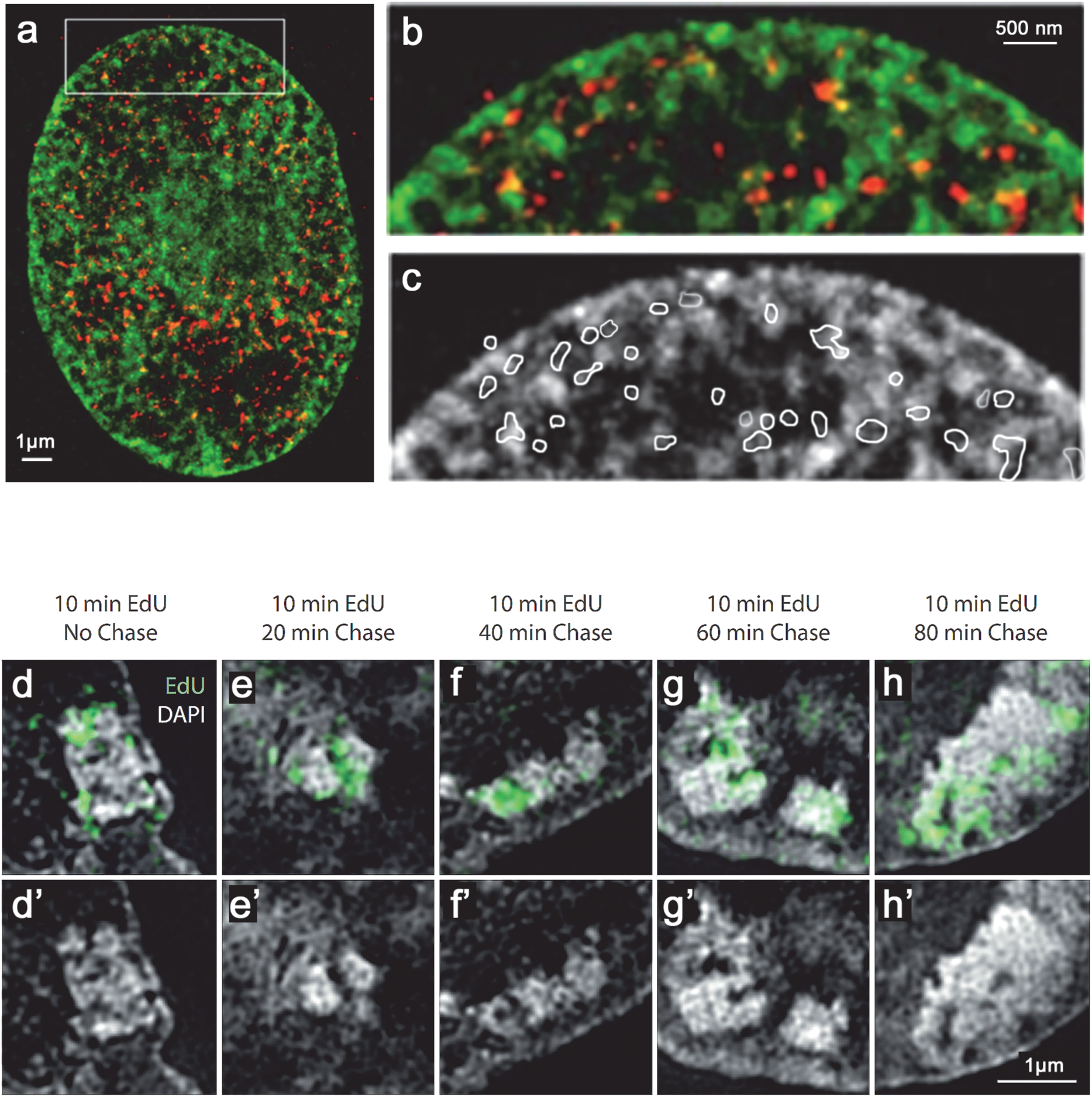
Evidence for DNA replication within the ANC and subsequent relocation of replicated DNA into the INC of heterochromatin blocks. a-c. Replication domains (RDs, red) in a HeLa cell nucleus at early S-phase, recorded with SMLM. The cell was fixed immediately after a 15 min pulse-labeling with the thymidine DNA base analog EdU, coupled with Alexa Fluor 555. The nuclear section demonstrates the location of RDs with newly replicated DNA (red) in the ANC, in particular on the surface of chromatin clusters (DNA stained with Vybrant Violet, green). **a** Dual color SMLM image of the entire nuclear section. **b-c** Magnification of boxed area shown in (a). **c** White outlines demonstrate the location of most EdU-labeled RDs at the periphery of chromatin domains (white-grey), expanding in section areas (black) attributed to the interchromatin compartment (IC). **d-h** EdU-pulse-chase experiment of heterochromatic blocks in undifferentiated mouse embryonic stem cells in mid-late S-phase, recorded with SIM. Optical sections show an overlay of DAPI stained DNA (white-gray) and replicated DNA after application of a 10min pulse with EdU (coupled to Alexa488; green). **d** Cell fixed immediately after the EdU-pulse demonstrates replicated DNA located mostly within low density DNA of wide IC-channels (attributed to the ANC). **e-h** cells fixed after increasing chase-time of 20/40/60/80 min before fixation show a relocation of labeled DNA into dense DNA of heterochromatic blocks (attributed to the INC). **d’-h’** DAPI stained DNA only. This result provides a glimpse into the dynamics of chromatin movements between immediately adjacent ANC and INC compartments.

**Supplemental Figure S8.**
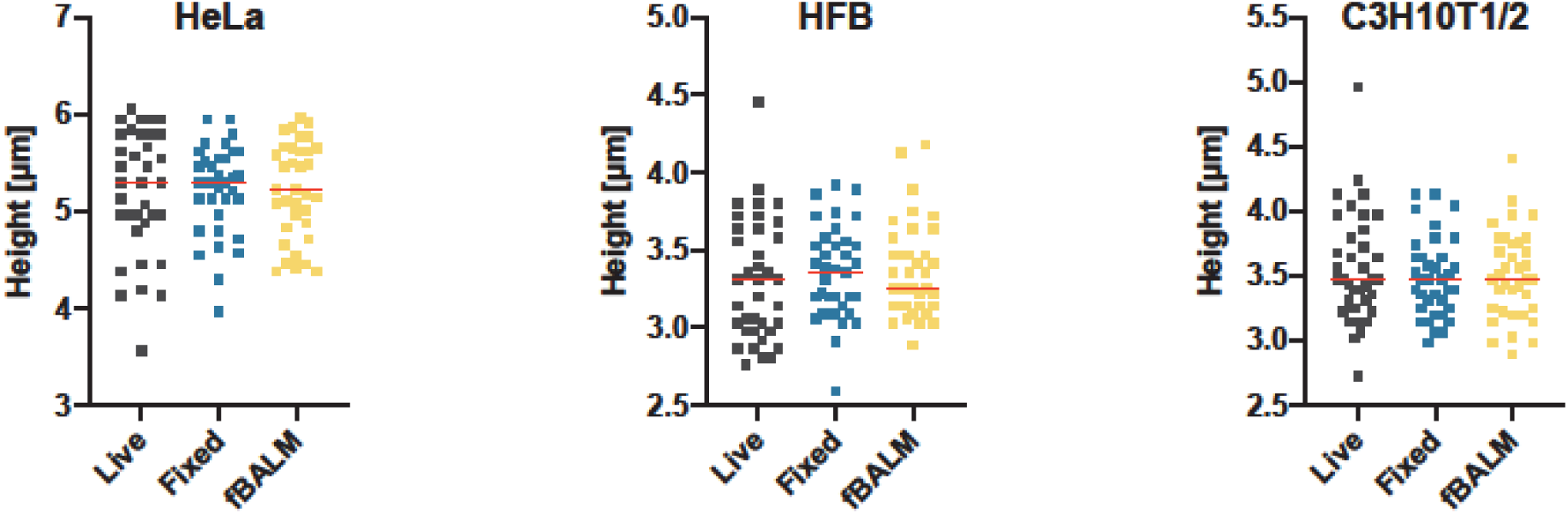
Measurements of cell-heights in live-cells, fixed cells in 1x dPBS and in fBALM imaging buffer. To test, whether cell fixation and high-resolution imaging with SMLM-fBALM has a flattening effect on nuclear morphology, we determined the height of HeLa nuclei (**a**), C3H10T1/2 (**b**) nuclei and HFB nuclei (**c**) in living and fixed cells, as well as in cells kept in fBALM imaging buffer. 20 min after adding Hoechst 33342, z-stacks with 200 nm intervals were recorded on a Visitron Spinning Disk microscope equipped with a 405 nm diode laser and a Photometrics Prime BSI camera using a 60x Nikon water immersion objective lens NA 1.2 attached to a Nikon Ti2 Microscopic stand. Nuclear heights were measured in image stacks with ImageJ in the x,y or y,z view. Red lines indicate the mean. Note that nuclear heights were not corrected for the limited depth resolution of the objective lens. Since this bias was the same for all recorded cells, it did not affect the comparative measurements. Conclusion: For each cell type, the nuclear heights were not significantly affected in living and fixed cells with or without incubation with fBALM buffer: HeLa (N=35, ANOVA P=0.98), HFb (N=37, ANOVA P=0.73) and C3H10T1/2 (N=36, ANOVA P=0.65).

**Supplemental Videos S1a and S1b**

Movie with the positions of blinking signals (SMPs) recorded from DNA bound Sytox Orange molecules (SMPs) in the BJ1 nucleus shown in Figure 2. The SM positions of a mid-plane optical section (thickness ∼ 600 nm) of this BJ1-nucleus with about 10 million localizations were visualized by assigning each SM position to a 10 nm sized pixel (corresponding to the average localization precision of 8.5 nm). Video S1a starts with the visualization of the entire nuclear section and then zooms in to smaller regions of interest (boxed area 2). Video S1b provides a view on the rotating 600 nm thick section.

**Supplemental Video S2**

Movie of a C3H 10T1/2 nucleus recorded for 6 hours after microinjection of three 20 nm sized nanobeads. Magnified images of two nanobeads in the ‘lower’ part of the nucleus are shown in Figure 5f,g. Other nanobeads or clusters were located in the cytoplasm, as demonstrated by serial sections. The display on the left shows the moving, intranuclear nanobeads. Notably, movements of nanobeads in the cytoplasm were more rapid and extensive. The right display demonstrates a random-walk like trajectories of intranuclear nanobeads (compare Figure 5h).

